# Ventral CA1 Population Codes for Anxiety

**DOI:** 10.1101/2023.09.25.559358

**Authors:** Sean C. Lim, Stefano Fusi, René Hen

## Abstract

The ventral hippocampus is a critical node in the distributed brain network that controls anxiety. Using miniature microscopy and calcium imaging, we recorded ventral CA1 (vCA1) neurons in freely moving mice as they explored variants of classic behavioral assays for anxiety. Unsupervised behavioral segmentation revealed clusters of behavioral motifs that corresponded to exploratory and vigilance-like states. We discovered multiple vCA1 population codes that represented the anxiogenic features of the environment, such as bright light and openness, as well as the moment-to-moment anxiety state of the animals. These population codes possessed distinct generalization properties: neural representations of anxiogenic features were different for open field and elevated plus/zero maze tasks, while neural representations of moment-to-moment anxiety state were similar across both experimental contexts. Our results suggest that anxiety is not tied to the aversive compartments of these mazes but is rather defined by a behavioral state and its corresponding population code that generalizes across environments.

## INTRODUCTION

Emotions are central internal states that are evoked by specific stimuli and organize perception, cognition, behavior, and physiology to produce an adaptive response^1,2^. Anxiety is an emotional state that is elicited by distal or potential threats and enables the animal to avoid harm enhancing its chances of survival. States of anxiety are characterized by heightened arousal and vigilance with associated patterns of risk assessment, defensive behaviors and autonomic responses depending on the environmental context and specific nature of the threat^3,4^. In humans, anxiety is often expressed as subjective tension and worried thoughts as well as physiological changes like increased heart rate, sweating, and dizziness. These cognitive and somatic effects typically promote vigilance and avoidance behaviors. Excessive, persistent, and disruptive anxiety states are clinically diagnosed as anxiety disorders, and while the precipitating stimuli are different depending on the disorder, all interfere with normal functioning and are subjectively distressing^5^.

To assess anxiety-like behavior in rodent models, tasks involving approach-avoidance conflict are often used. These tasks are thought to measure the degree to which an animal internally balances the drive to explore a novel area and the desire to avoid danger in exposed or brightly illuminated regions of the maze^6^. Many of these behavioral paradigms, including the open field test, elevated plus maze (EPM), and elevated zero maze (EZM), have been validated as assays of anxiety-like behavior using anxiolytic pharmacology^7–10^.

Seminal studies have shown that the hippocampus is essential for cognitive processes including episodic memory and spatial navigation^11–13^. However, the hippocampus is also extremely sensitive to stress^14,15^ and has been implicated in human affective disorders, including anxiety^16^ and depression^17^. Furthermore, it is becoming increasingly recognized that the dorsal and ventral hippocampus possess distinct functional specializations^18–20^. Chronic lesions of the ventral hippocampus are anxiolytic and produce altered processing of innately aversive stimuli including predator odor^21^, open areas^22^, and footshocks^23^. Furthermore, acute and reversible optogenetic manipulation of the ventral hippocampus and its projections directly affect anxiety-associated avoidance behavior^24,25^.

Recent studies have begun to elucidate how the ventral hippocampus is involved in the encoding of anxiogenic stimuli. Using single cell analyses, they have demonstrated that some neurons in ventral CA1 exhibit increased activity during the exploration of the exposed open arms of the EPM^24,26^. However, it is possible that several important features that characterize anxiety states can be detected only when the analysis is extended to populations of neurons, because the information is distributed across multiple cells^27^. In particular, it would be important to understand if there exist features of the neural representations of anxiety that are preserved across different environments and tasks. Their existence would indicate that in vCA1 there is an abstract representation of anxiety that would allow a downstream readout to report the anxiety state in multiple situations. Interestingly, these representations might still encode the specific sensory features of the situations which the animal experiences. These types of abstract representations have been observed in multiple brain areas for a variety of different cognitive variables^28^.

Here, we utilized *in vivo* freely moving calcium imaging in mice in combination with unsupervised behavioral segmentation and neural population analysis to investigate how vCA1 encodes anxiogenic features of the environment and anxiety-like behavioral states.

## RESULTS

### High Lux Decreases Exploration and Increases Avoidance Behavior

We used virally expressed GCaMP7f and miniature microendoscopy to perform calcium imaging in vCA1 of freely-moving mice while they explored a sequence of environments. A gradient refractive index (GRIN) lens was implanted over vCA1 allowing for visualization of Ca^2+^ transients in individual neurons expressing virally delivered GCaMP7f (Figure 1A, 1B, 1C)^29^. To process our data, we utilized a recently developed cell segmentation algorithm, constrained non-negative matrix factorization for microendoscopy (CNMF-E)^30,31^.

**Figure 1.**
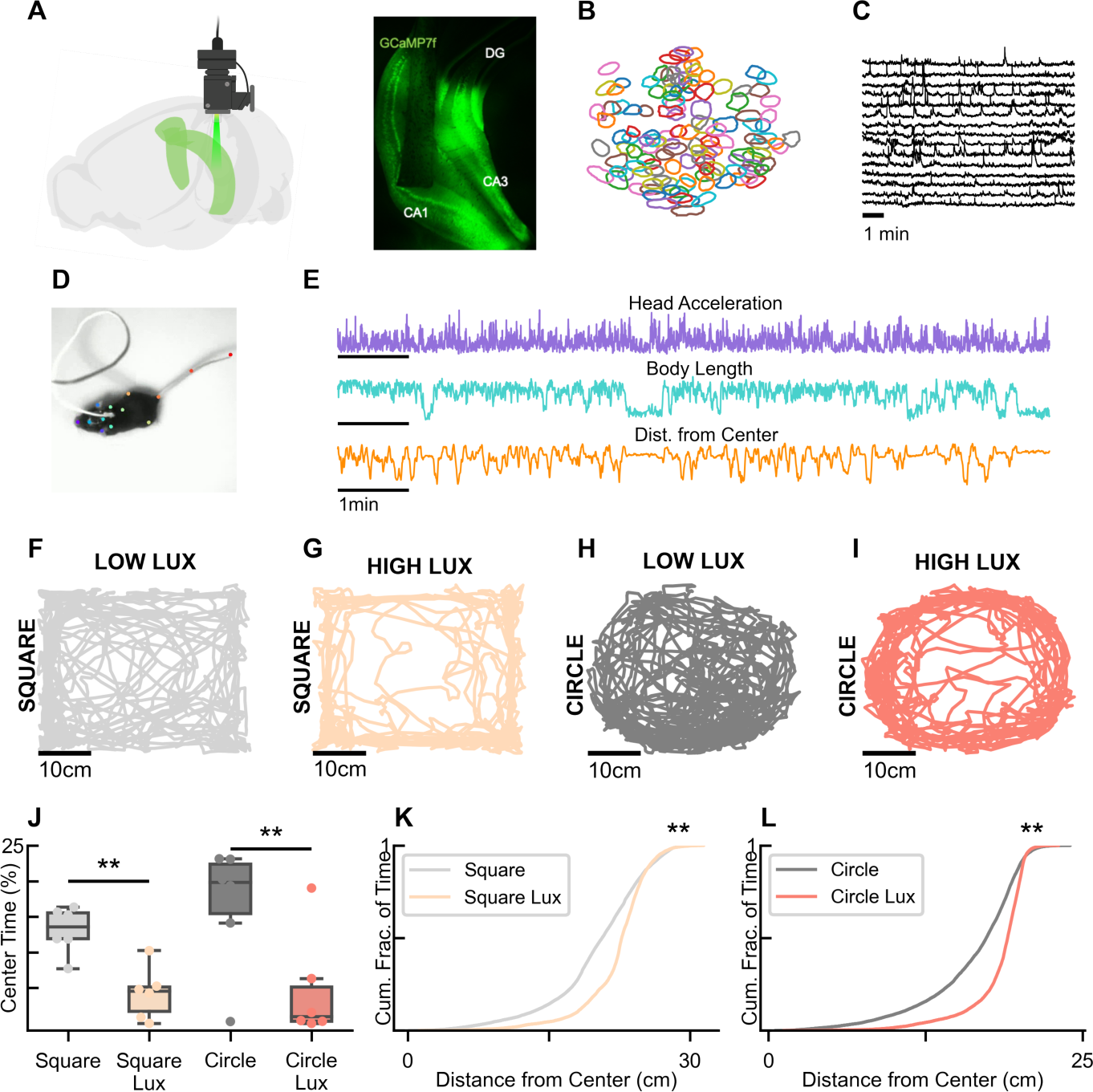
High lux increases avoidance behavior in the open field. **(A)** Summary of experimental setup: GCaMP7f is expressed in vCA1 neurons *in vivo* and imaged with a gradient refractive index (GRIN) lens and miniaturized microscope. Confocal image of vCA1, showing GCaMP7f-expressing neurons and the lens tract after removal of the GRIN lens **(B)** An example field-of-view from one mouse showing the spatial footprints of CNMF-E identified neurons. **(C)** Example calcium transients from 15 neurons from the same mouse during a single behavioral session. **(D)** A single frame showing the subject mouse and body parts labeled with DeepLabCut. **(E)** Example time series traces of head acceleration, body length, and distance from the center for a single mouse. **(F)** An example location tracking trace for a single mouse during the low lux, square open field session. **(G)** An example location tracking trace for a single mouse during the high lux, square open field session. **(H)** An example location tracking trace for a single mouse during the low lux, circle open field session. **(I)** An example location tracking trace for a single mouse during the high lux, circle open field session. The time spent in the center region of the open field arenas is significantly decreased during high lux sessions compared to low lux session (Two-way ANOVA significant main effect of lux, F_(1,20)_= 17.043, p < 0.01). **(K)** The distribution of distances from the center in the square arena are significantly shifted away from the center during high lux sessions compared to low lux sessions (Kolmogorov-Smirnov test, D_(6)_= 0.278, p < 0.01). **(L)** The distribution of distances from the center in the circle arena are significantly shifted away from the center during high lux sessions compared to low lux sessions (Kolmogorov-Smirnov test, D_(6)_= 0.320, p < 0.01).

Simultaneously captured behavioral data was acquired using an IR sensitive camera enabling dissociation of experimental lighting conditions from video recording. Additionally, the miniature microscopes which were used to record neural activity also enabled collection of three-axis head acceleration data (Figure 1E). To reliably track specific body points on the mice in the behavioral videos we used DeepLabCut, a newly developed pose-estimation package (Figure 1D)^32^.

Experimental modulation of the light intensity (lux) above an open field arena is a robust means to alter the exploratory and avoidance behaviors exhibited by mice^33^. We utilized square and circle open field arenas in combination with high and low lux levels to experimentally manipulate the physical environment and the putative anxiety level of the mice. In the high lux conditions, we found a significant decrease in the percent time exploring the center of the arenas (Figure 1F, 1G, 1H, 1I, 1J) as well as a smaller number of center entries, less percent distance travelled in the center and less distance travelled overall (Figure S1A, S1B, S1C). Furthermore, by looking at the distance from the center (Figure 1E), a continuous metric that does not require discretizing the arena into a center region, we found that during the high lux conditions mice spent significantly more time further away from the center (Figure 1K, 1L, S1D, S1E, S1F, S1H, S1I, S1J). These results suggest that the high lux manipulation induces an internal state which drives increased avoidance and decreased exploratory behavior, consistent with a state of increased anxiety.

Two additional behavioral tasks that are often used to investigate anxiety-like behavior in rodents are the EPM and EZM. Both tasks involve an internal conflict between exploration and avoidance of novel but exposed and elevated regions of the mazes. We utilized a series of EPM and EZM tasks with the open arms facing different spatial locations of the room to assess exploratory and anxiety-like behaviors while controlling for visual cues in different parts of the larger experimental context. In each of the EPM and EZM sessions we observed significantly lower exploration of the open arms, suggesting that these regions are innately aversive (Figure S1J, S1K).

### Unsupervised Behavioral Segmentation Reveals Motifs that are Modulated by Anxiogenic Features of the Environment

Traditional metrics for the analysis of the open field, such as percent-time-in-center and number-of-center-entries, have been well validated using anxiolytic pharmacology but collapse behavioral data to a handful of values. To capture more of the behavioral variability that occurs during each open field session we utilized an autoregressive-hidden-Markov-model-based unsupervised behavioral segmentation^34,35^ approach to identify behavioral motifs using head accelerometer and body length data acquired from the miniature microscope and DeepLabCut tracking respectively (Figure 2A). With this method, we found six behavioral motifs that occurred across the sequence of open field sessions (Figure 2B). These motifs exhibited distinct distributions of head acceleration and body length (Figure 2C, 2D, 2E, 2F, 2G, 2H). The frequency of occurrence for each motif was significantly modulated by the high versus low lux conditions with only a few motifs showing significant differences across the square versus circle arena shapes (Figure 2I, 2J). Furthermore, a linear support vector machine (SVM) could distinguish when an individual animal was in the high lux or low lux conditions using only the frequencies of these motifs across time without access to any information about center occupancy (Figure 2K). However, these motifs did not allow a linear decoder to distinguish the square versus circle arena shapes (Figure 2K). Several motifs were visually identified to correspond with known behaviors such as freezing (m3), investigation (m2), and scanning (m1)^36^ (Supplementary Video). However, not every motif had a clearly identifiable visual appearance. This might be due to the limited ability of our top-down camera angle to capture minute changes in head movements and body posture. Notably, the motifs that were increased by the high lux manipulation included those which visually corresponded to defensive and vigilance-like behaviors, while the motifs that were decreased included those which corresponded to exploratory-like behaviors. Additionally, the position and center occupancy of the animals were not used for the unsupervised behavioral segmentation suggesting that the behavioral motifs extracted are providing additional information about the behavioral state of the animal that is related to but distinct from the traditional metrics.

**Figure 2.**
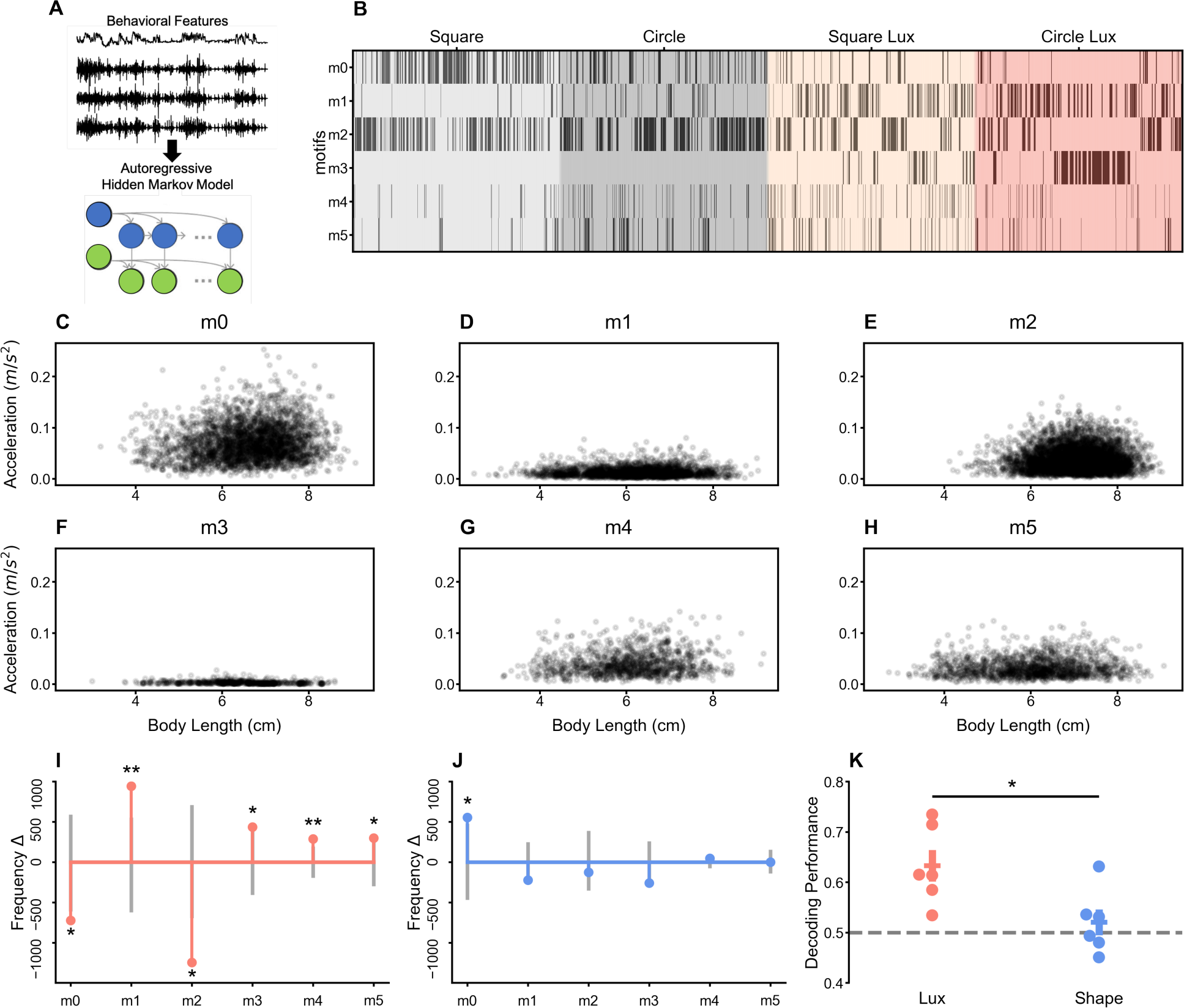
Unsupervised behavioral segmentation reveals motifs that distinguish high and low lux conditions. **(A)** Diagram of procedure for unsupervised behavioral segmentation. **(B)** An example motif raster plot showing the occurrence of each behavioral motif (m0 to m5) from a single mouse across the four open field sessions. **(C)** A scatterplot showing the relationship between body length and total head acceleration during motif m0. **(D)** A scatterplot showing the relationship between body length and total head acceleration during motif m1 for all mice. **(E)** A scatterplot showing the relationship between body length and total head acceleration during motif m2 for all mice. **(F)** A scatterplot showing the relationship between body length and total head acceleration during motif m3 for all mice. **(G)** A scatterplot showing the relationship between body length and total head acceleration during motif m4 for all mice. **(H)** A scatterplot showing the relationship between body length and total head acceleration during motif m5 for all mice. **(I)** All motif frequencies are significantly different between high and low lux sessions. **(J)** Only motif m0 frequencies are significantly different between square and circle sessions. The observed frequency differences are represented by the colored bars and points. Grey bars represent the 95^th^ and 5^th^ percentile values of the null distribution. Asterisks represent the significance of motif frequency differences where * is p < 0.05 and ** is p < 0.01. **(K)** A linear SVM trained on behavioral motifs can distinguish between high and low lux sessions (t-test, t_(5)_= 3.82, p < 0.05) but not square and circle arena shapes (t-test, t_(5)_= 0.99, p = 0.37). Lux decoding performance is significantly higher than shape decoding performance (paired t-test, t_(5)_= 3.51, p < 0.05).

Using a similar statistical approach, we extracted an independent set of behavioral motifs from the EPM and EZM sessions. Like in the open field experiments, these motifs varied in frequency across the length of each session (Figure S2A) and exhibited distinct acceleration and body length profiles (Figure S2B, S2C, S2D, S2E, S2F, S2G). Furthermore, these motifs were performed with different frequencies in the open and closed regions of the EPM and EZM (Figure S2I), and allowed a linear SVM to distinguish when an individual animal was in the open or closed regions without direct access to information about zone occupancy (Figure S2H).

In both sequences of experiments, groups of behavioral motifs were modulated by anxiogenic features of the environment suggesting that they reflect an underlying anxiety-like state.

### vCA1 Population Activity Distinguishes Lux Level in the Open Field

We computed the selectivity of each recorded vCA1 neuron for high versus low lux as well as for square versus circle arena shapes. Roughly 2/3 of the neurons exhibited significantly elevated activity in either the high or low lux conditions, with a similar proportion significantly modulated by the square or circle arena shapes (Figure 3A). Many of these neurons exhibited an increased frequency of calcium transients in the conditions specified (Figure 3B).

**Figure 3.**
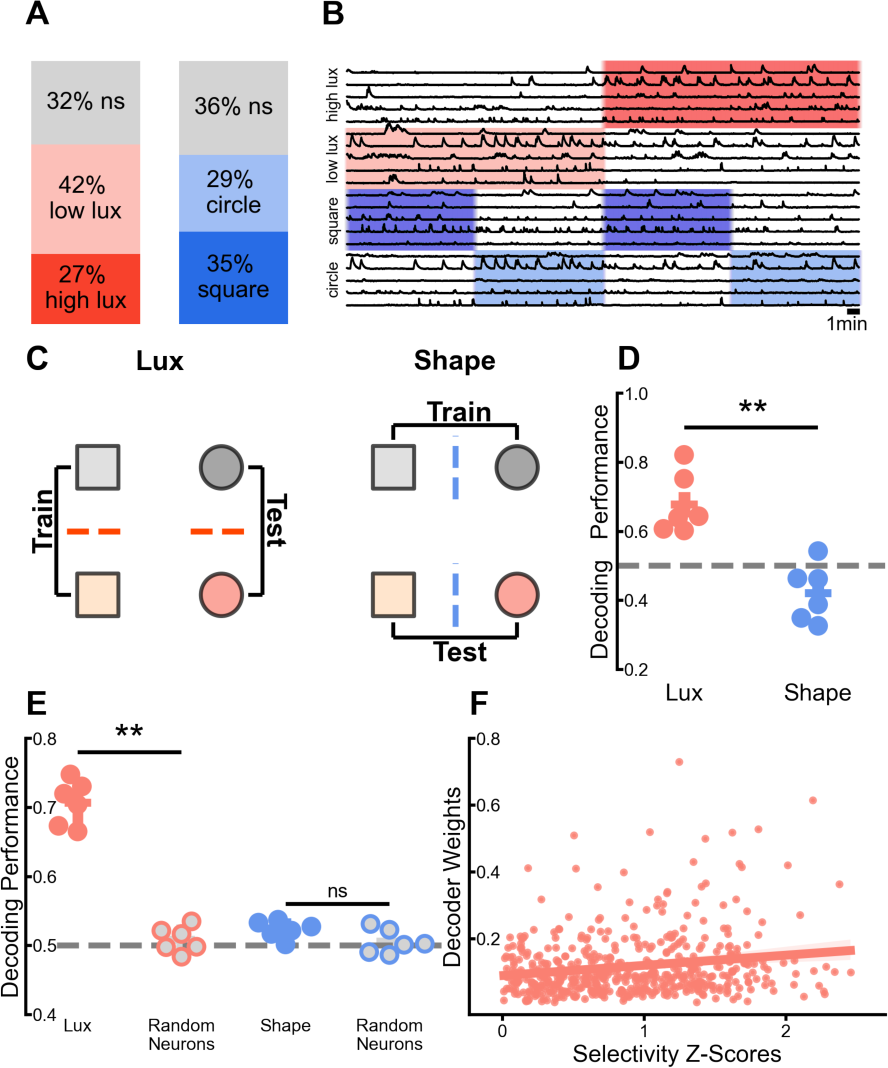
A vCA1 neural population code represents lux level. **(A)** Relative proportions of single neurons that are activated by high lux sessions (27%), activated by low lux sessions (42%), lux non-selective (32%), activated by square sessions (35%), activated by circle sessions (29%), or shape non-selective (36%). **(B)** Example calcium transients for high lux, low lux, square, and circle activated neurons across all four open field sessions. **(C)** Diagram depicting procedure performed for decoding high vs. low lux and square vs. circle sessions from vCA1 neural data. **(D)** Lux decoding performance is significantly above chance levels (paired t-test, t_(5)_= 7.38, p < 0.01); shape decoding performance is not significantly above chance levels (paired t-test, t_(5)_= −1.97, p = 0.11); lux decoding performance is significantly higher than shape decoding performance (paired t-test, t_(5)_= 4.29, p < 0.01). **(E)** Cross-mouse lux decoding performance is significantly higher using CCA aligned neural data compared to randomly selected neurons (paired t-test, t_(5)_= 12.06, p < 0.01); cross-mouse shape decoding performance is not significantly different using CCA aligned neural data compared to randomly selected neurons (paired t-test, t_(5)_= 1.88, p = 0.22). **(F)** A linear regression between selectivity z-scores and lux decoder weights for all recorded neurons is significant but with small magnitude (linear regression, R^2^= 0.03, F_(1,500)_= 16.92, p < 0.01).

We also determined the ability of the neural population to represent high versus low lux by using a linear SVM decoder. To assess whether the vCA1 representations of lux level allowed for generalization across arena shape, we trained the decoder on one pair of high and low lux conditions and tested its classification ability on a different pair with overall performance being averaged across all train and test pairs (Figure 3C)^28^. We found that the performance of the lux decoders was significantly above chance. Additionally, we performed a similar analysis in which we decoded the arena shape, training only on one pair of conditions. The arena shape decoders did not perform at above chance levels (Figure 3D). We reasoned that the ability to decode lux level from the vCA1 neural population could be due to differences in other variables, such as behavioral motif frequency or distance from the center, that were influencing vCA1 activity independently of lux. To account for these potential influences on vCA1 decoding ability we sampled balanced amounts of neural activity occurring during each motif and distance from the center in each session and repeated the lux decoding procedure (Figure S3A). Strikingly, we found that even with neural activity samples that were balanced for motifs and distance from the center (Figure S3B), the decoder could still classify high versus low lux across conditions (Figure S3C).

We next determined if the degree of high or low lux selectivity for single neurons identified previously was related to importance for the population level decoding performance. To assess this, we took the decoder weights for each neuron, which are a measure of how important each cell is for the performance of the decoder, and then performed a linear regression with the high and low lux selectivity z-scores, which indicate how strongly the activity of a neuron is modulated by the lux level. This revealed a significant but weak positive correlation between a neuron’s importance for decoding and degree of selectivity (Figure 3F). We also performed a complementary analysis where we removed a small number of neurons that were lux selective or non-selective and repeated the lux decoding procedure assessing the magnitude of the drop in decoding performance compared to the full neural population (Figure S3D). We found that there was a significant but small difference in decoding performance drop when removing lux selective versus non-selective neurons (Figure S3E). These results suggest that there is a weak relationship between single cell selectivity and contribution to the vCA1 neural population representation of lux level, but this does not exclude the possibility of selective neurons influencing the population code through mechanisms such as pairwise or higher-order correlations.

To determine if the neural representations of lux could generalize across different mice we sampled equal periods of neural activity from each of the high and low lux sessions in pairs of mice, and then aligned the neural activity from each mouse using canonical correlation analysis (CCA)^37,38^. This procedure allowed us to train a decoder to classify lux level using data from one mouse and test its performance on data from another mouse, with overall performance averaged across all pairs of mice (Figure S3F). In contrast to a selection of random neurons from each mouse, the CCA-aligned neural activity allowed for cross-mouse lux decoding at levels significantly above chance (Figure 3E). A similar analysis performed for shape decoding produced a much lower performance which was essentially indistinguishable from decoders trained with unaligned random neurons (Figure 3E).

Because one of the primary metrics used to evaluate avoidance behavior in the open field is center occupancy time, we wondered whether vCA1 also encoded distance from the center. We binned the continuous distance from the center data into 5 bins across all the open field sessions and trained a decoder to distinguish between different distance bins, with a chance decoding level of approximately 0.2 (Figure S4A). Using this approach, we found that decoding performance was above chance at discriminating distance from the center (Figure S4B).

However, we also performed an analogous set of control analyses where we trained a decoder to distinguish ‘north’ versus ‘south’ or ‘east’ versus ‘west, again using 5 bins for each analysis, (Figure S4A) and observed that the decoding performance was also above chance levels (Figure S4B). Notably, the distance from the center decoding performance was not significantly different from the north-south or east-west decoding performances suggesting that the representation of distance from the center may be more related to the encoding of spatial information rather than a component of an anxiety-like state. These results demonstrate that vCA1 encodes multiple types of information simultaneously and importantly, implies that the abstract representation of lux level (i.e. decoding across arena shape) is not a trivial consequence of only a single variable being encoded in the network.

### vCA1 Population Activity Distinguishes Open and Closed Regions of the EPM and EZM

For the EPM and EZM data, we computed single neuron selectivity for open versus closed regions across all the sessions. Like previous studies of vCA1 in the EPM^24^, we found that approximately 1/2 of the neurons exhibited significantly elevated activity in the open regions with a smaller proportion demonstrating significantly increased activity in the closed regions (Figure 4A, 4B).

**Figure 4.**
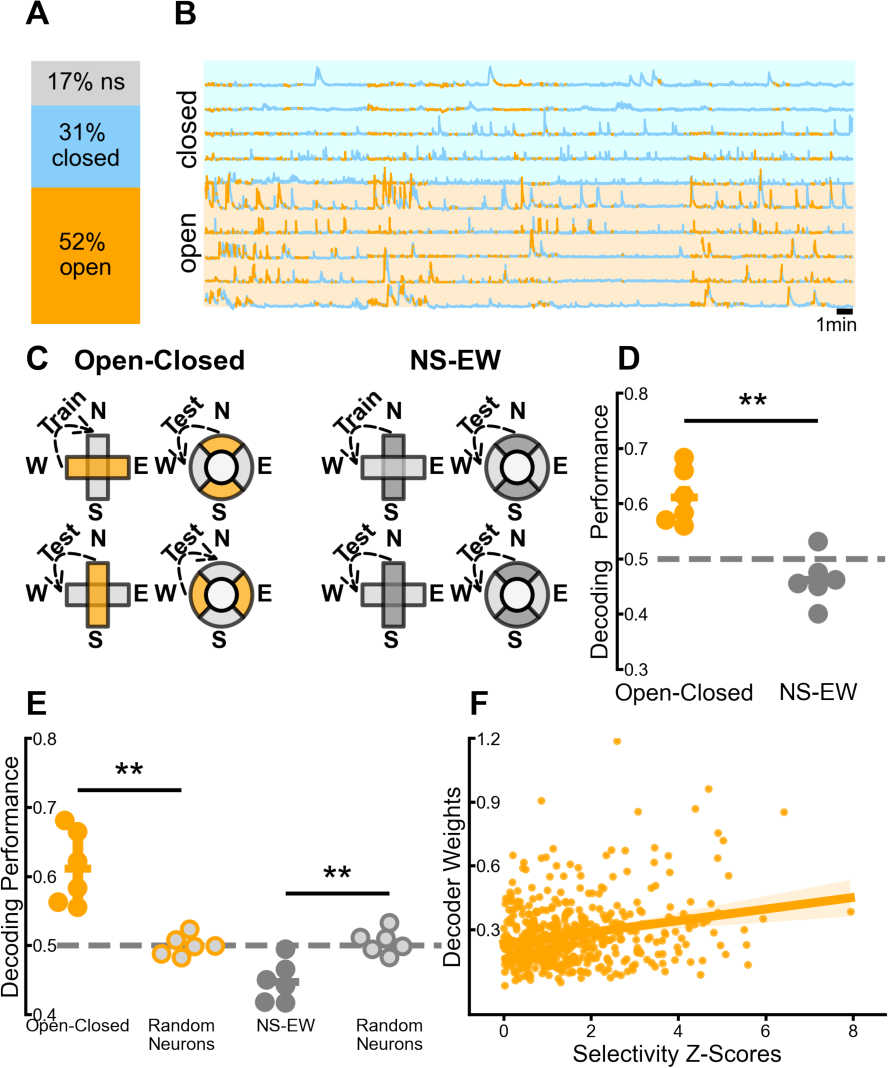
A vCA1 neural population code represents open and closed regions. **(A)** Relative proportions of single neurons that are activated by open regions (52%), activated by closed regions (31%), or region non-selective (17%). **(B)** Example calcium transients for open and closed activated neurons across all four elevated plus and zero maze sessions. Orange segments of the trace are time periods when the mouse is in the open regions. Blue segments of the trace are time periods when the mouse is in the closed regions. **(C)** Diagram depicting procedure performed for decoding open vs. closed regions and NS vs. EW regions from vCA1 neural data. **(D)** Open-closed decoding performance is significantly above chance levels (paired t-test, t_(5)_= 8.18, p < 0.01); NS-EW decoding performance is not significantly different from chance levels (paired t-test, t_(5)_= −2.58, p = 0.05); open-closed decoding performance is significantly higher than NS-EW decoding performance (paired t-test, t_(5)_= 5.75, p < 0.01). **(E)** Cross-mouse open-closed decoding performance is significantly higher using CCA aligned neural data compared to randomly selected neurons (paired t-test, t_(5)_= 6.13, p < 0.01); cross-mouse NS-EW decoding performance is significantly lower using CCA aligned neural data compared to randomly selected neurons (paired t-test, t_(5)_= −3.72, p < 0.05). **(F)** A linear regression between selectivity z-scores and open-closed decoder weights for all recorded neurons is significant but with small magnitude (linear regression, R^2^= 0.06, F_(1,499)_= 29.67, p < 0.01).

Additionally, we used decoding analysis to determine if the neural population dynamics encoded open versus closed regions. To assess whether vCA1 representations generalized across the arena shapes, we trained the decoder to distinguish open versus closed regions on one EPM or EZM session and tested its classification ability on a completely different session with the overall decoding performance averaged across all train and test pairs. (Figure 4C). We found that the decoding performance for distinguishing open versus closed regions was significantly above chance (Figure 4D). In an analogous set of control analyses, we trained decoders to distinguish the north and south regions of the arenas from the east and west regions (Figure 4C), and found that the decoding performances were not above chance levels (Figure 4D). Furthermore, it was not possible to decode either the arena shape or the orientation of the arena (Figure S5C, S5D). Using a motif balanced decoding analysis like the one described above for the open field sessions (Figure S5A), we found that even when neural samples were balanced for motifs, the decoder was still able to classify open versus closed regions across conditions (Figure S5B).

We then assessed the relationship between open and closed region selectivity and importance for population decoding by comparing the decoder weights to the open and closed selectivity z-scores. This analysis, like those described above for high and low lux, demonstrated a significant but slight positive correlation between importance for decoding and degree of selectivity (Figure 4F).

Subsequently, we applied a cross-mouse decoding analysis, like the one describe above for the open field data, to determine if the neural representations of open and closed regions could generalize across different mice. We trained a decoder to distinguish open versus closed regions using data from one mouse and tested its performance on data from another mouse averaging the performance across all pairs of mice (Figure S3F). CCA aligned neural activity but not unaligned random neurons enabled cross-mouse decoding of open versus closed regions at levels significantly above chance (Figure 4E). Additionally, a comparable procedure performed for north-south versus east-west decoding revealed performance which was significantly below that of decoders trained with random neurons (Figure 4E).

### vCA1 Population Activity Distinguishes Motif Clusters in the Open Field

To investigate the neural representations of behavioral motifs in vCA1, we generated population vectors that were composed of the average firing rate for each neuron during the specified motif. For each animal, there was a single population vector for each behavioral motif. We computed the Pearson’s correlation coefficient for each pair of population vectors in each animal, providing us a measure of how similar the neural representations of all the behavioral motifs were to each other. The average pattern of correlations between these neural population vectors revealed two clusters of motifs (Cluster 1: m0, m2; Cluster 2: m1, m3, m4, m5) (Figure 5B). Interestingly, these clusters corresponded to the groups of motifs that were modulated by high lux in an opposing fashion (Figure 2I). We wondered if these motif clusters were related to the traditional metrics of avoidance in the open field, so we performed a pair of linear regression analyses between the frequency of each motif cluster and the percent center exploration time during each session. Our results demonstrated that the frequency of motif cluster 1 was strongly correlated with center exploration (Figure 5D) while the frequency of motif cluster 2 was strongly anti-correlated with center exploration time (Figure 5C). Intriguingly, two of the motifs in cluster 2 were visually identified as freezing (m3) and scanning (m1) behaviors, while one of the motifs in cluster 1 appeared to be investigation (m2), as described above (Supplementary Video). These findings suggest that these motif clusters might correspond to exploratory and vigilance-like behavioral states which appear to be directly represented at the neural population level in vCA1. Additionally, the marked correlation and anti-correlation of the motif clusters to center exploration as well as their prominent modulation by the high lux manipulation (Figure 5A) implies that they may be reflecting an underlying anxiety state.

**Figure 5.**
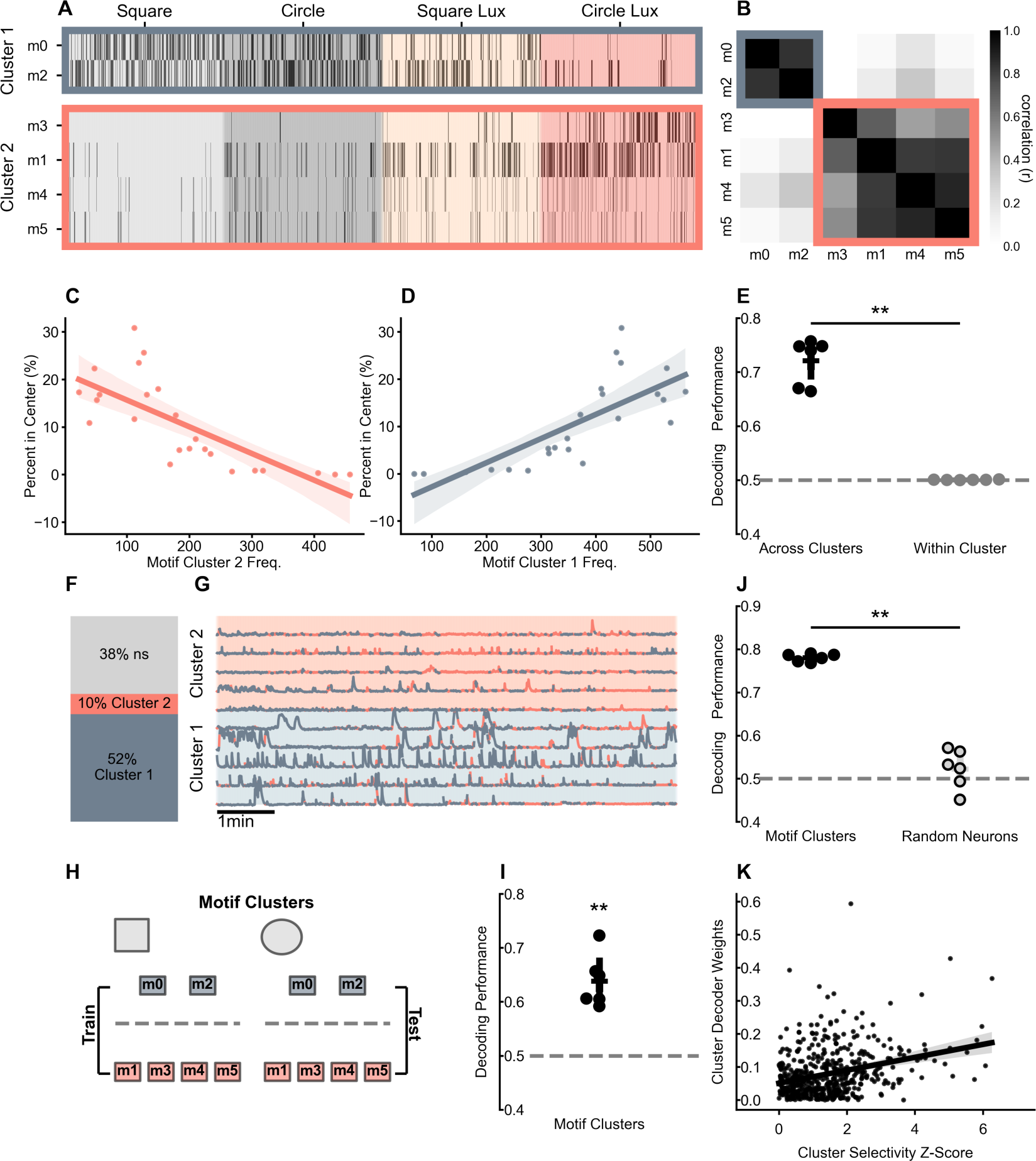
A vCA1 neural population code represents behavioral motif clusters in the open field. **(A)** An example motif raster plot showing the occurrence of each behavioral motif (m0 to m5) from a single mouse across the four open field sessions grouped by motifs in Cluster 1 (grey) and motifs in Cluster 2 (salmon). **(B)** Population vector correlation coefficient matrix ordered to show the neural similarity between Cluster 1 (exploration) and Cluster 2 (vigilance) motifs. **(C)** A linear regression between the frequency of Cluster 2 motifs and the percent time spent in the center; each point is a single behavioral session from a single mouse (linear regression, R^2^= 0.56, F_(1,22)_= 28.17, p < 0.01). **(D)** A linear regression between the frequency of Cluster 1 motifs and the percent time spent in the center; each point is a single behavioral session from a single mouse (linear regression, R^2^= 0.58, F_(1,22)_= 29.88, p < 0.01). **(E)** Across cluster decoding performance is significantly higher than chance levels (t-test, t_(5)_= 12.86, p < 0.01); within cluster decoding performance is not significantly different than chance levels (t-test, t_(5)_= 1.94, p = 0.11); across cluster decoding performance is significantly higher than within cluster decoding performance (paired t-test, t_(5)_= 12.91, p < 0.01). **(F)** Relative proportions of single neurons that are activated by Cluster 1 motifs (52%), activated by Cluster 2 motifs (10%), or cluster non-selective (38%). **(G)** Example calcium transients for Cluster 1 and Cluster 2 activated neurons across all four open field sessions. Salmon segments of the trace are time periods when the mouse is exhibiting a Cluster 2 motif. Grey segments of the trace are time periods when the mouse is in the closed regions. **(H)** Diagram depicting procedure performed for decoding Cluster 1 vs. Cluster 2 motifs from vCA1 neural data. **(I)** Cluster decoding performance is significantly above chance levels (paired t-test, t_(5)_= 5.91, p < 0.01). **(J)** Cross-mouse cluster decoding performance is significantly higher using CCA aligned neural data compared to randomly selected neurons (paired t-test, t_(5)_= 13.05, p < 0.01). **(K)** A linear regression between selectivity z-scores and cluster decoder weights for all recorded neurons is significant but with small magnitude (linear regression, R^2^= 0.09, F_(1,500)_= 50.66, p < 0.01).

These patterns of population vector correlations were observed in each individual animal not just in aggregate across all the mice (Figure S6C, S6E, S6G, S6I, S6K, S6M). In a complementary set of analyses, we trained a series of linear decoders to distinguish pairs of behavioral motifs from vCA1 neural activity (Figure S6A). High decoding performance would indicate that the neural representations of each behavioral motif are distinct, while low decoding performance would suggest that the neural representations are more similar and thus less distinguishable. The average level of decoding performance revealed two clusters of motifs which matched those from the population vector analysis (Figure S6B, S6D, S6F, S6H, S6J, S6L, S6N). The most significant difference was that the motif pair decoding showed that motif m3 (freezing) had a particularly unique neural representation, although it was less distinguishable from other cluster 2 motifs compared to the cluster 1 motifs.

To further probe these motif cluster representations, we trained a decoder to distinguish motif cluster 1 (exploration) and motif cluster 2 (vigilance) using neural data from a single pair of behavioral motifs and tested its performance on data from another pair of behavioral motifs that were not used for training. The averaged decoding performance for those classifiers was significantly above chance levels and markedly higher compared to decoders trained on pairs of motifs within the same cluster and tested on pairs of the opposite cluster (Figure 5E).

We wondered if these motif clusters would drive single neuron tuning in vCA1, so we computed the neural selectivity for motif cluster 1 versus motif cluster 2 for all recorded cells. Strikingly, approximately 1/2 of the neurons demonstrated significantly increased activity during motif cluster 1, with a much smaller proportion exhibiting elevated activity during motif cluster 2 (Figure 5F, 5G).

We also assessed the ability of the neural population to represent motif clusters by using decoding analysis. To evaluate whether the vCA1 representations of motif clusters allowed for generalization across arena shape, we trained the decoder to distinguish motif cluster 1 versus motif cluster 2 using data from either the square or circle sessions and tested its performance on data from the other arena, with overall performance being averaged across all train and test pairs (Figure 5H). We found that the decoding performances were significantly above chance suggesting that vCA1 encodes these motif clusters at the population level (Figure 5I).

We then evaluated the relationship between motif cluster selectivity and population decoding performance by comparing the decoder weights to the motif cluster selectivity z-scores. This analysis demonstrated a significant but small positive correlation between the decoder weights and degree of selectivity (Figure 5K). This was notable because the proportion of motif cluster selective neurons was quite divergent with more than 50% activated by cluster 1 and only ∼10% activated by cluster 2 (Figure 5F).

We used a cross-mouse decoding procedure to assess if the neural representations of motif clusters could generalize across mice, training a decoder to distinguish motif clusters using data from one mouse and testing its performance on data from another mouse with performance averaged across all pairs of mice (Figure S3F). Neural activity aligned with CCA but not unaligned random neurons enabled cross-mouse decoding of motif clusters significantly above chance levels (Figure 5J).

### vCA1 Population Activity Distinguishes Motif Clusters in the EPM and EZM

The vCA1 neural representations of behavioral motifs in the EPM and EZM data exhibited a remarkably similar pattern of correlation to those from the open field, with two clusters of motifs (Cluster 1: m2, m3; Cluster 2: m0, m1, m4, m5) (Figure 6B) that exhibited differential expression in the open and closed regions of the EPM and EZM (Figure S2I). Furthermore, we found that the frequency of motif cluster 1 was significantly correlated with open region exploration (Figure 6D) while the frequency of motif cluster 2 was significantly anti-correlated with open region exploration (Figure 6C) suggesting that these motif clusters might also correspond to exploratory and vigilance-like behavioral states directly encoded in vCA1 neural population activity. Notably, the cluster 1 motifs (exploration) are expressed both in the closed and open regions while the cluster 2 motifs (vigilance) are more highly expressed in the closed regions (Figure 6A, S2I). This implies that when the animals are within the open regions of the EPM and EZM they may be in a more exploratory-like behavioral state, challenging the more typical interpretation of EPM and EZM behavior which holds that the animals are in a state of higher anxiety within the open regions of the arenas. Like in the open field, similar population vector correlations were observed in each individual animal (Figure S7C, S7E, S7G, S7I, S7K, S7M) with motif pair decoding analysis (Figure S7A) revealing analogous clusters of motifs (Figure S7B, S7D, S7F, S7H, S7J, S7L, S7N), that even showed a motif (m4) that was more distinguishable than any other, like motif m3 (freezing) from the open field. Decoders trained to distinguish motif cluster 1 and motif cluster 2 from one pair of motifs and tested on a held-out pair of motifs exhibited performance that was significantly above chance and much higher than the performance of decoders trained on pairs of motifs within the same cluster and tested on pairs of the opposite cluster (Figure 6E).

**Figure 6.**
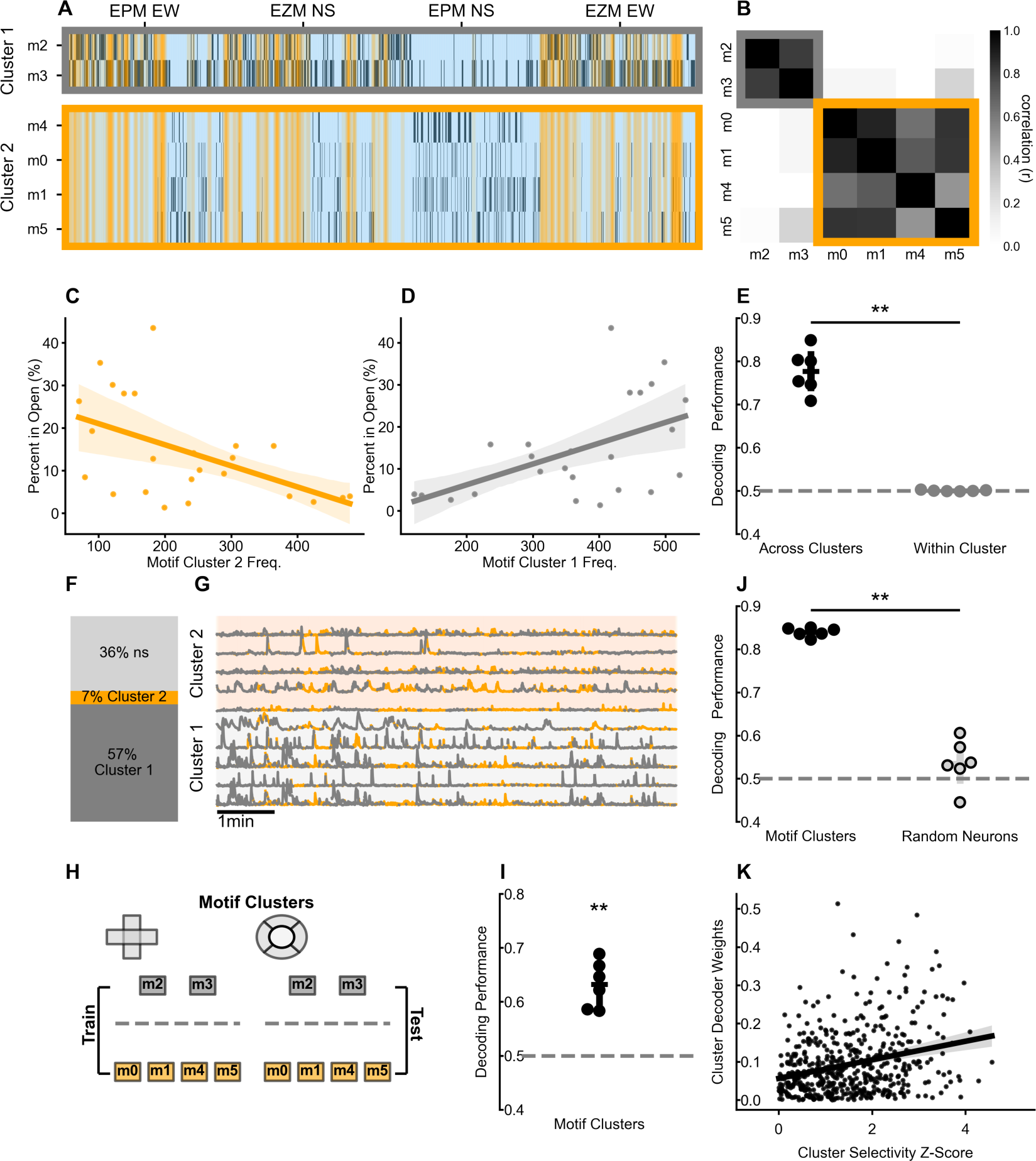
A vCA1 neural population code represents behavioral motif clusters in the elevated plus/zero mazes. **(A)** An example motif raster plot showing the occurrence of each behavioral motif (m0 to m5) from a single mouse across the four elevated plus and zero maze sessions grouped by motifs in Cluster 1 (grey) and motifs in Cluster 2 (orange). **(B)** Population vector correlation coefficient matrix ordered to show the neural similarity between Cluster 1 (exploration) and Cluster 2 (vigilance) motifs. **(C)** A linear regression between the frequency of Cluster 2 motifs and the percent time spent in the open regions; each point is a single behavioral session from a single mouse (linear regression, R^2^= 0.27, F_(1,22)_= 7.976, p < 0.01). **(D)** A linear regression between the frequency of Cluster 1 motifs and the percent time spent in the open regions; each point is a single behavioral session from a single mouse (linear regression, R^2^= 0.27, F_(1,22)_= 7.976, p < 0.01). **(E)** Across cluster decoding performance is significantly higher than chance levels (t-test, t_(5)_= 13.51, p < 0.01); within cluster decoding performance is not significantly different than chance levels (t-test, t_(5)_= 0.51, p = 0.63); across cluster decoding performance is significantly higher than within cluster decoding performance (paired t-test, t_(5)_= 13.67, p < 0.01). **(F)** Relative proportions of single neurons that are activated by Cluster 1 motifs (52%), activated by Cluster 2 motifs (10%), or cluster non-selective (38%). **(G)** Example calcium transients for Cluster 1 and Cluster 2 activated neurons across all four elevated plus and zero maze sessions. Orange segments of the trace are time periods when the mouse is exhibiting a Cluster 2 motif. Grey segments of the trace are time periods when the mouse is in the closed regions. **(H)** Diagram depicting procedure performed for decoding Cluster 1 vs. Cluster 2 motifs from vCA1 neural data. **(I)** Cluster decoding performance is significantly above chance levels (paired t-test, t_(5)_= 8.62, p < 0.01). **(J)** Cross-mouse cluster decoding performance is significantly higher using CCA aligned neural data compared to randomly selected neurons (paired t-test, t_(5)_= 15.72, p < 0.01). A linear regression between selectivity z-scores and cluster decoder weights for all recorded neurons is significant but with small magnitude (linear regression, R^2^= 0.08, F_(1,499)_= 43.24, p < 0.01).

Like in the open field, more than 1/2 of the neurons had significantly elevated activity during motif cluster 1, with a much smaller proportion showing increased activity during motif cluster 2 (Figure 6F, 6G). Additionally, decoders trained to discriminate motif cluster 1 (exploration) from motif cluster 2 (vigilance) from either the EPM or EZM sessions and tested on data from the held-out arena (Figure 6H) exhibited performances that were significantly above chance, suggesting that vCA1 encodes these motif clusters at the population level in a format that generalizes across arenas (Figure 6I). Linear regression analysis also revealed a significant but weak positive correlation between the decoder weights and motif cluster selectivity z-scores, like the analyses described above (Figure 6K). Furthermore, cross-mouse decoding (Figure S3F) showed that neural activity aligned through CCA but not unaligned random neurons enabled the generalization of motif cluster representations, with above chance decoding performance (Figure 6J).

### vCA1 Representations of Anxiogenic Features of the Environment and Motif Clusters Have Distinct Generalization Properties

We next determined if the representation of lux and the representation of the open regions were encoded in a fashion which allowed for generalization of the aversive stimuli (i.e. high lux and open regions) across the open field, EPM, and EZM tasks. This kind of format would imply that all anxiogenic features of the environment should produce a similar population response in vCA1, or should be encoded in the same subspace of neural activity. To directly assess this hypothesis, we trained a decoder to distinguish high versus low lux conditions in the open field assays and tested its performance at distinguishing the open versus closed regions of the EPM and EZM; overall performance was averaged with the converse train-test groups (Figure 7A).

**Figure 7.**
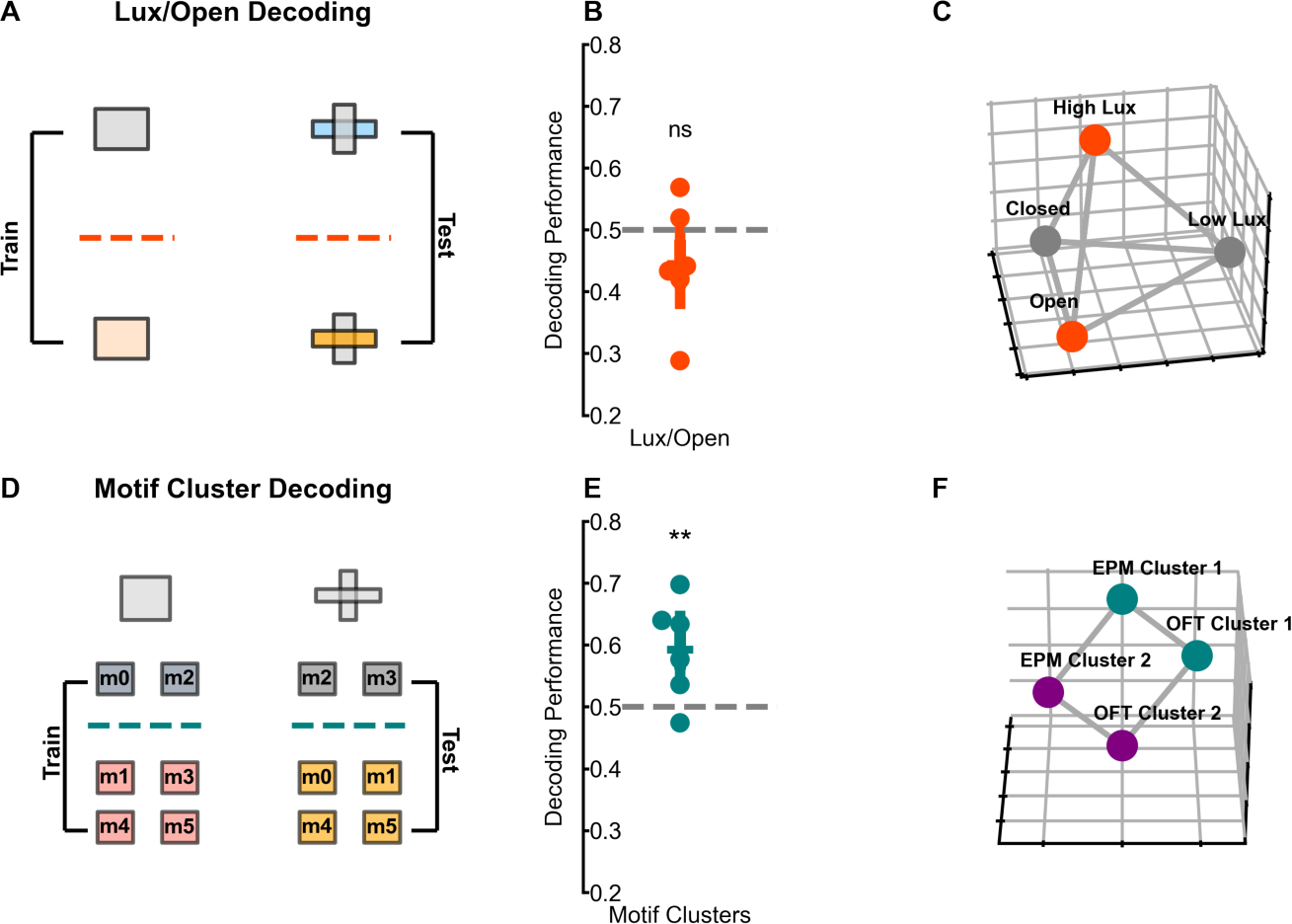
vCA1 representations of behavioral motif clusters generalize across open field and elevated plus/zero maze sessions. **(A)** Diagram depicting procedure performed for decoding high lux/open regions vs. low lux/closed regions from vCA1 neural data. **(B)** Decoding performance is not significantly different from chance levels (paired t-test, t_(5)_= −0.27, p = 0.79). **(C)** Diagram depicting neural population activity geometry that does not allow for decoding of anxiogenic features across experimental contexts. **(D)** Diagram depicting procedure performed for decoding Cluster 1 and Cluster 2 motifs from vCA1 neural data. **(E)** Decoding performance is significantly above chance levels (paired t-test, t_(5)_= 8.84, p < 0.05). **(F)** Diagram depicting neural population activity geometry that allows for motif cluster decoding across experimental contexts.

Intriguingly, we observed that the decoder performance was not above chance (Figure 7B). This suggests that the vCA1 neural population code for anxiogenic features of the environment maintains the information about feature identity and does not collapse the representation so that all anxiogenic features become indistinguishable (Figure 7C) and implies that they are not encoded within the same subspace. The underlying representational geometry in the activity space is similar to the one depicted in Figure 7C, where 4 independent clusters represent the 4 possible conditions (high/low lux, open/closed regions). The centroids of these clusters define a high-dimensional tetrahedron. Notice that the coding direction for high/low lux is approximately orthogonal to the coding direction for discriminating between open and closed regions.

We also wanted to determine if the vCA1 representations of motif clusters would generalize across the open field, EPM and EZM. Using a similar procedure as described above, we trained a decoder to discriminate between motif cluster 1 (exploration) and motif cluster 2 (vigilance) in the open field and tested its performance at discriminating motif cluster 1 and motif cluster 2 in the EPM and EZM (Figure 7D). Importantly, the behavioral motifs in the open field and EPM/EZM were independently extracted, meaning that they are not identical movements.

Despite this, we found that the decoder performance was above chance levels (Figure 7E), implying that vCA1 encodes these motif clusters in a format which allows the representations to generalize across markedly distinct contexts (Figure 7F). This generalized neural representation of exploratory and vigilance-like motifs aligns with the idea that anxiety is an emotional state which can evoke similar behaviors in different contexts.

## DISCUSSION

### vCA1 Represents Anxiogenic Features of the Environment Through a Stimulus-Specific Population Code

Previous studies have suggested that vCA1 carries a representation of innately anxiogenic features of the environment through specialized cells with characteristic projection targets that categorically encode the intensity of the animals’ anxiety state^24^. Indeed, our work here confirms that vCA1 encodes anxiogenic features of the open field, EPM, and EZM. However, we find that the relationship between selectivity and importance for population decoding is weak. This is analogous to results from work that has investigated spatial representations in dorsal CA1 and DG and found that position is not encoded only by classically defined place cells^27^. We also find that vCA1 encoding of anxiogenic features allows for generalization across arena shapes, but not across different kinds of features (i.e. high lux and open regions are not encoded within the same subspace). This suggests that vCA1 neural representations of anxiogenic features are stimulus-specific and may directly encode differences in sensory quality (e.g. lux level versus ‘open-ness’). In other words, the anxiety state might be similar in the high lux and open maze situations, but its neural representation in vCA1 is not disentangled from the particular sensory characteristics of each anxiogenic feature^28^. Importantly, the EPM and EZM tasks were performed in low light conditions, enhancing the differences between the anxiogenic stimuli in the open field and the elevated mazes. Stimulus-specific representations of aversive stimuli have been found in animal as well as human neural imaging data^39,40^, and these kinds of representations may be part of the neural substrate which drives the production of different adaptive responses in distinct aversive conditions. Our work here suggests that vCA1 encodes aversive features through a population code that enables the separation of stimuli with divergent sensory properties. Additional *in vivo* recording experiments in vCA1 using a larger variety of stimuli will be necessary to determine if this hypothesis holds true within and across sensory modalities.

### vCA1 Represents Exploratory and Vigilance-like States Through an Abstract Population Code

Functional manipulations of vCA1 cell bodies and projections have demonstrated that this brain region is causally involved in driving avoidance^24,25^. However, previous studies have largely focused on coarse, region-based behavioral metrics. Here, we utilized unsupervised behavioral segmentation to characterize moment-to-moment behavioral motifs as animals explored well-validated tasks used to study anxiety-like behavior. In contrast to other brain regions such as the dorsal striatum^41,42^, vCA1 does not seem to encode each behavioral motif independently but rather as two larger clusters. These behavioral states roughly correspond to exploratory and vigilance-like behaviors, similar to recent studies of the amygdala^43,44^. Importantly, this representation generalizes across open field and elevated plus/zero maze tasks, in contrast to the vCA1 population encoding of anxiogenic features which is stimulus-specific. Indeed, these representations of behavioral states are consistent with the encoding of an underlying anxiety state in vCA1. This suggests that the behavioral motif clusters extracted here may better capture moments of anxiety than prior region-of-interest approaches. Recent studies have focused on the importance of the ventral hippocampus for mediating approach-avoidance conflict^45,46^, and our findings support this idea, demonstrating a neural representation of exploration and vigilance-like behavioral states in vCA1. Furthermore, we show that vigilance-like behaviors are not limited to periods when the animals are directly experiencing an aversive stimulus, like when mice freeze in the closed regions of the EPM and EZM, which suggests that these states have some degree of persistence. Moreover, our results show that mice can exhibit anxiety-related behaviors even when they are not within the center of the open field or the open arms of the EPM. Capturing these previously unmeasured moments of vigilance and their associated neural population responses allows us to define a neural activity subspace that may reflect the moment-to-moment anxiety level of the animals and could serve as a neurophysiological endophenotype^47^ for preclinical animal models of anxiety disorders^48,49^. Further work is necessary to investigate whether these anxiety state representations extend to other aversive situations and whether they are modulated by anxiolytic pharmacological treatments.

### Multiple vCA1 Population Codes Enable Simultaneous Encoding of Anxiety State and Anxiogenic Sensory Stimuli

In both the open field and elevated plus/zero maze tasks vCA1 simultaneously encodes the relevant anxiogenic features of the environment as well as the moment-to-moment anxiety state of the animal. Numerous studies have linked vCA1 to a variety of different functions including anxiety^24–26,50^, reward^51,52^, fear^53^, learning^54,55^, and social behavior^56–58^. Our results suggest that vCA1 may play a role in such a diversity of behaviors through multiple population codes which exist in different subspaces of neural activity^43,59^. This assertion challenges current models of vCA1 function which posit that neurons with different projection targets selectively encode different facets of stimuli and experience. A few seminal studies have directly investigated the activity of vCA1 neurons with defined projections and observed that there are differences in single neuron selectivity during a variety of behavioral tasks^24,26^. Our results show that the coordinated and distributed dynamics of the vCA1 neural population allow for an additional layer of information representation that is distinct from single cell activity.

Population coding may thus be a fundamental property of vCA1 that allows for the simultaneous encoding of moment-to-moment internal anxiety states as well as external anxiogenic sensory stimuli in complex environments.

## ACKNOWLEDGEMENTS

We thank the current and past members of the Hen Lab including Jack Berry, Jessica Jimenez, Clay Lacefield, Wei-Li Chang, Gergely Turi, and Phi Nguyen for useful comments and discussion. We also thank Steve Siegelbaum and Mark Ansorge for helpful comments on this work and scientific discussion. This work was funded by the NIMH Ruth L. Kirschstein F30 Fellowship MH124424 (S.C.L), and the Hope for Depression Research Foundation and NIMH R01 MH068542 (R.H.).

## AUTHOR CONTRIBUTIONS

Conceptualization, S.C.L, S.F., R.H.; methodology, S.C.L.; software, S.C.L.; formal analysis, S.C.L.; investigation, S.C.L.; data curation, S.C.L.; writing – original draft, S.C.L.; writing – review & editing, S.C.L., S.F., R.H.; visualization, S.C.L.; supervision, S.F., R.H.; funding acquisition, S.C.L., R.H.

## METHODS

### Experimental Models and Subject Details Animal Subjects

All procedures were conducted in accordance with the U.S. NIH Guide for the Care and Use of Laboratory Animals and the New York State Psychiatric Institute Institutional Animal Care and Use Committees at Columbia University. 8-week old adult male C57BL/6J mice were supplied by Jackson Laboratory. Mice were housed singly and kept on a 12-hour light-dark cycle in air-filtered, temperature- and humidity-controlled conditions with ad libitum food and water access; experiments were conducted during the light phase.

### Viral Constructs

For calcium imaging, viruses (AAV9-syn-jGCaMP7f-WPRE-SV40) were packaged by Addgene at titers of approximately 1 x 10^13^ vg/mL. Viral aliquots were diluted to approximately 3 x 10^12^ vg/mL in distilled water (ddH_2_O).

## Method Details

### Stereotactic Surgeries Viral Injection

For *in vivo* calcium imaging in vCA1, subject mice were given 5 mg/kg carprofen analgesic and anesthetized with 2-5% isoflurane. Fully anesthetized animals were maintained at 1.5% isoflurane with an oxygen flowrate of 1L/min and placed in a stereotactic frame (David Kopf, Tujunga, CA). Eyes were lubricated with an ophthalmic ointment, and body temperature was maintained at 37°C with a warm water circulator (Stryker, Kalamazoo, MI). Prior to the initiation of surgical procedures, fur was shaved, the incision site sterilized with alternating betadine and 70% ethanol scrubs, and a craniotomy was drilled at the vCA1 coordinates (AP −3.16mm, ML +3.5mm, DV −3.85, −3.5, −3.25mm). Dura was removed from the craniotomy site using curved forceps (Fine Science Tools, Foster City, CA) and the site was irrigated with sterile saline (NaCl 0.9%) to prevent coagulation. A glass pipette containing the virus was lowered to the desired depth. A Nanoject II (Drummond Scientific, Broomall, PA) was used to inject 32.2nL of virus at intervals of 10 seconds to obtain the desired volume. One minute was allowed to pass between injections at different z-axis levels. Following the completion of microinjections, five minutes were allowed to pass before the pipette was retracted from the brain tissue. Viral injection coordinates are in mm with Bregma as reference.

### Gradient Index (GRIN) Lens Implantation

Immediately following viral injection, an additional surgical procedure was performed while the animal was maintained at 1.5% isoflurane anesthesia. Three skull screws (Antrin Miniature Specialties, Fallbrook, CA) were inserted in evenly spaced locations around the craniotomy site. An integrated GRIN lens and baseplate (Inscopix, Palo Alto, CA: 0.5 mm diameter, 6.1 mm length) was slowly lowered in 0.1 mm increments to a depth of −3.7 mm and fixed to the skull with dental cement (Parkell, Brentwood, NY). Additional subcutaneous saline and carprofen were provided perioperatively and for three days postoperatively to prevent dehydration and provide analgesia. Animals were observed postoperatively until they regained righting reflex and ambulated normally. Imaging experiments did not begin until at least three weeks after surgery to allow time for recovery and sufficient viral expression.

### Behavioral Assays

Behavioral assays were conducted under low light conditions unless otherwise specified and all experiments were recorded using Ethovision XT 11.5 (Noldus, Leesburg, VA) with infrared sensitive monochromatic digital cameras (Basler, Ahresberg, Germany).

#### Open field

Mice experienced four 10-minute sessions of open field exploration within a single day. In between behavioral sessions, mice were removed from the arena and placed into a transfer cage with bedding. The arenas were cleaned with 70% ethanol before each session. The sequence of sessions was conducted in the same order for each mouse starting with a square shaped arena (40 x 40 x 30cm length-width-height) followed by a circular arena created using an opaque plastic circular insert (40cm diameter) placed inside the original open field box. These were followed by sessions using the identical square and circle arena but with a bright light (∼600 lux) positioned over the center of the arena.

#### Elevated plus/zero mazes

Mice experienced four 10-minute sessions using either a standard configuration elevated plus maze (13.5’’ height of the maze from the floor, 25’’ full length of each arm-type, 2’’ arm width, 7’’ tall closed arm walls, with 0.5’’ tall/wide ledges on the open arms) or elevated zero maze (55cm diameter with 2’’ width, 5’’ tall closed region walls, with 0.5’’ tall ledges on the open regions) with the open regions of each maze facing either east and west or north and south. All elevated plus and elevated zero maze sessions were conducted with ambient light levels less than 10 lux. In between behavioral sessions, mice were removed from the arena and placed into a transfer cage with bedding. The arenas were cleaned with 70% ethanol before each session. The sequence of sessions was conducted in the same order for each mouse starting with the elevated plus maze with an east-west orientation, followed by the elevated zero maze in a north-south orientation. This was followed by the elevated plus maze in a north-south orientation and the elevated zero maze in an east-west orientation.

### Behavioral Data Processing

#### Pose estimation

Behavioral videos were analyzed using the DeepLabCut software package^32^. A neural network was trained to track eleven mouse body parts: nose, right ear, left ear, head, neck, right shoulder, left shoulder, body, tail base, tail mid, and tail tip. These body parts were manually labelled on a set of ∼50-100 still frames from each behavior video and from these labeled examples the network was trained to detect the specified body parts. Once the neural network was sufficiently trained (train root mean square error < 5 pixels), the tracking data was used for subsequent analysis with custom Python scripts.

#### Definition of regions-of-interest and occupancy calculation

For every behavioral video acquired, a single frame was extracted and the napari image viewing package was used to draw regions-of-interest over the image. Furthermore, a scale line was drawn on the image to enable conversion of pixel tracking and distances to centimeters. In the open field a single region-of-interest was drawn which encompassed the precise floor of the specific arena. This region was used to compute the center point of both the square and circle arenas. The center point was subsequently used to define center regions of either 20cm length for the square arena or 20cm diameter for the circle arena. Additionally, this center point was used to define a wall region which encompassed the region between a 35cm length square or 35cm diameter circle and the edge of the arena. In the elevated plus and elevated zero mazes two lines were manually drawn on the images which bisected each other dividing the arenas into open and closed regions. In this way, the center region of the elevated plus maze was partitioned into four equal regions which were considered part of the closest open or closed arm. Regions-of-interest were visually verified before proceeding with analysis. Tracking data acquired from DeepLabCut processing was used in combination with the defined regions-of-interest to determine the occupancy of each body part in all the regions at a given moment in time, producing a Boolean time-series for each combination. For the region occupancy data reported above, only the head body part was considered.

#### Distance from the center calculation

The distance from the center was calculated for all the open field assays by using the previously defined center point and computing the Euclidean distance between the head of the animal and the center point for each time point.

#### Body length calculation

The body length of the animal was calculated by taking the Euclidean distance between the head and the tail base of the animal for each time point.

#### Accelerometer processing

Raw accelerometer data was acquired from the Inscopix nVista3.0 miniature microscope (Inscopix, Palo Alto, CA) by using the Inscopix Data Acquisition software (Inscopix, Palo Alto, CA) and exported to CSV using the Inscopix Data Processing software (Inscopix, Palo Alto, CA). Three-axis (x, y, z) acceleration time series were resampled depending on the downstream analysis.

### Unsupervised Behavioral Segmentation

The *ssm* package^60^ was used to fit an autoregressive hidden Markov model (ARHMM) to the three-axis accelerometer and body length data. As described in Wiltschko et al. 2015 and Batty et al. 2019, an ARHMM consists of an HMM, which models the behavioral features provided as a sequence of discrete states and transition probabilities, in combination with an autoregressive observation model, which captures stereotyped time varying dynamics within these states. The three-axis acceleration and body length data for each animal were first resampled to 1 Hz for compatibility with subsequent decoding analyses. The data was then centered and scaled to unit variance and concatenated forming one continuous set of behavioral features for all the animals recorded. A first-order ARHMM was fit to these behavioral features enabling the detection of motifs across animals.

#### Number of motifs selection

To determine an estimate for the number of states that best fits the data a ten-fold cross validation approach was used. A series of ARHMMs was fit to a subset of the behavioral data while varying the number of states used with each iteration. The log-likelihood of the held-out data was evaluated for each ARHMM and averaged across the ten train/test splits. These values were used as guide to select the number of states for downstream analyses.

### Calcium Imaging Data Acquisition and Processing

Before each imaging session, mice were anesthetized briefly to attach the nVista3.0 miniature microscope (Inscopix, Palo Alto, CA) to the baseplate, after which mice recovered for 30min in a transfer cage with bedding before beginning the imaging and behavioral experiments. Calcium videos were recorded from the miniature microscopes with the Inscopix Data Acquisition software (Inscopix, Palo Alto, CA). Behavior and calcium imaging videos were synchronized with a TTL-triggering system (a TTL pulse from Ethovision XT 11.5 and Noldus IO box was received by the nVista3.0 data acquisition box to begin each recording session). Calcium videos were acquired at 20 frames-per-second and the LED power settings were calibrated for each mouse and maintained across sessions.

#### Calcium imaging data preprocessing, motion correction and cell segmentation

All raw calcium videos were preprocessed in the same manner using the Inscopix Data Processing software (Inscopix, Palo Alto, CA). Videos were temporally downsampled to 5 frames-per-second and spatially downsampled by a binning factor of 4. Calcium videos were motion corrected using the Python implementation of NoRMCorre^61^ in the *CaImAn* software package^31^. Cells were segmented from the calcium videos using the Python implementation of Constrained Non-negative Matrix Factorization for microEndoscopic data (CNMF-E)^62^ in the *CaImAn* software package^31^. Neurons were verified by manual inspection, rejecting any cells that did not have appropriate spatial footprints and calcium temporal dynamics. CNMF-E produces three ouputs: the raw calcium trace (C_raw_), the deconvolved trace (C), and the deconvolved spikes (S), which provides an estimate of the number of spikes that occurred in each time bin. The estimated noise (s_n_) was calculated using the formula *s*_n_ = *c*_raw_– *c*. A minimum spike size (s_min_) of *s*_min_ = 15 × *s*_n_ was applied to remove several tiny estimated spikes from the calcium trace. All three traces were normalized by the standard deviation of the noise (s_n_). For all subsequent downstream analyses C_raw_ was used exclusively. Analyses which focus on either open field or elevated plus/zero maze use calcium transient data segmented exclusively from calcium videos corresponding to the specific sequence of experiments. For analyses that encompassed both open field and elevated plus/zero maze experiments, calcium videos were concatenated across both sequences of experiments to allow for segmentation of common cells across experimental days.

### Quantification and Statistical Analysis

#### Behavioral metrics

Custom Python scripts were used to compute the percent time in zone, number of entries, percent distance travelled in zone, and total distance travelled in each behavioral session.

#### Distance from center distributions

Distances from the center were assessed as described above. These data were combined for all the animals to determine the overall distribution of distances from the center for each type of session, which were represented using empirical cumulative distribution functions.

#### Motif frequency

Motif frequency differences were calculated by aggregating the motif frequencies for all the animals in each condition and taking the difference between the overall frequencies. Null distributions of motif frequency differences were generated by shuffling the motifs across time for each animal and recalculating the overall frequency differences.

#### Linear regressions between motif cluster frequency and behavioral metrics

Motif clusters were initially defined based on the groups of motifs which were positively or negatively modulated by the high lux manipulation in the open field. For the sequence of elevated plus/zero maze experiments, motif clusters were grouped based on feature space similarity to the previously defined open field motifs.

#### Using motifs to predict lux and open regions

For analyses using motifs to predict whether animals were experiencing high lux in the open field or exploring open regions of the elevated plus/zero mazes, a linear SVM was used for classification and the target variable was balanced so there were equal instances of each decoded class.

#### Cell selectivity analysis

Neurons were classified as selective for a particular pair of behavioral variables (e.g. open versus closed regions of the elevated plus/zero mazes, center versus wall regions of the open field assays) as previously described^63^. Briefly, the calcium activity traces were shuffled 1000 times and the mean rate was computed for time spent during one of the behaviors and subtracted from the mean rate during time spent in the other specified behavior, generating a null distribution of activity rate differences for each neuron. The actual (non-shuffled) mean rate difference was compared to the shuffle distribution using a nonparametric two-sided permutation test. Rate difference z-scores were calculated for each neuron in relation to the null distribution. Neurons with rate difference z-scores that were one standard deviation above or below the mean of the null distribution were deemed selective for that behavior. A neuron that did not reach this threshold for either behavior was considered non-selective.

#### Population decoding analysis

For all decoding analyses neural data was downsampled to 1 Hz as previously described^27^. Decoding was performed with custom Python scripts using the LinearSVC module in the *scikit-learn* package^64^. For analyses involving two-class discrimination, neural data was limited to periods when one of these two classes occurred. In cases where decoded variables were unbalanced due to natural differences in behavior, neural data was sampled to balance the occurrence of the target classes. A linear SVM decoder was trained to classify a specific variable dichotomy (e.g. high lux versus low lux) from one subset of conditions (e.g. square arenas) and then the performance of the decoder was tested on a completely held-out subset of conditions (e.g. circle arenas). The performance of the decoder for distinguishing this dichotomy was determined for multiple train and test conditions and averaged across these groups to estimate the decoding performance for that specific variable.

To estimate a null distribution for decoding analyses, a similar procedure was used as previously described^28,65^. Briefly, the decoding performance was kept intact by performing four independent random solid rotations of the neural activity space. For each condition, the neuron indices were shuffled to rotate the patterns corresponding to that condition in the neural activity space. After this rotation, a null-model value for decoding performance was computed as described above. This procedure was repeated twenty times to obtain a null distribution of decoding performance values.

#### Motif and distance from the center balanced decoding analyses

Motif balanced decoding analyses were performed by evenly sampling neural data occurring during each motif while simultaneously keeping the samples of decoded variables equal. For certain analyses, distance from the center was also balanced in a similar fashion.

#### Canonical correlation analysis and cross-mouse decoding

Canonical correlation analysis (CCA) was used to align the neural dynamics from two animals so that cross-animal decoding could be performed^37,38^. Canonical correlation analysis attempts to find patterns that are common across two data sets by finding latent factors, which are linear combinations of the individual units in each population, such that the factors from each data set are maximally correlated. Because each animal freely explores the arenas it is exposed to, the same time points do not necessarily correspond to similar behaviors or region-of-interest occupancies across animals. To address this, a defined number of samples was taken for each variable from each mouse. In this way, samples from each animal were curated so that they corresponded behaviorally at equivalent time points. CCA was then applied to these curated samples from two animals to align the appropriately corresponding neural data. The number of latent factors is a user defined parameter; analyses described above used 30 factors, but similar results were obtained using a reasonable range of factors. Using these aligned data sets, decoding analysis was performed as described above for every pairwise grouping of animals, with decoding performance averaged across all pairs of mice. As a control analysis, curated samples were obtained from 30 random neurons selected from each mouse, with decoding analysis performed identically.

#### Selective cell ablation decoding analysis

To assess the relative importance of selective neurons for decoding performance a small subset (ten neurons) of either selective or non-selective neurons were removed from the population before decoding analysis was performed. The difference in decoding performance using the entire population of cells and the performance with the subset removed was calculated for one hundred repetitions and compared between selective and non-selective groups.

#### Linear regression between selectivity and decoder weights

Linear regressions were performed between the selectivity z-scores calculated for each neuron during the selectivity analysis and the weights assigned to each neuron by the decoder during population decoding analysis. The decoder weights used here were the average weights for each train/test pair in the specified decoding analysis.

---

**Figure S1.**
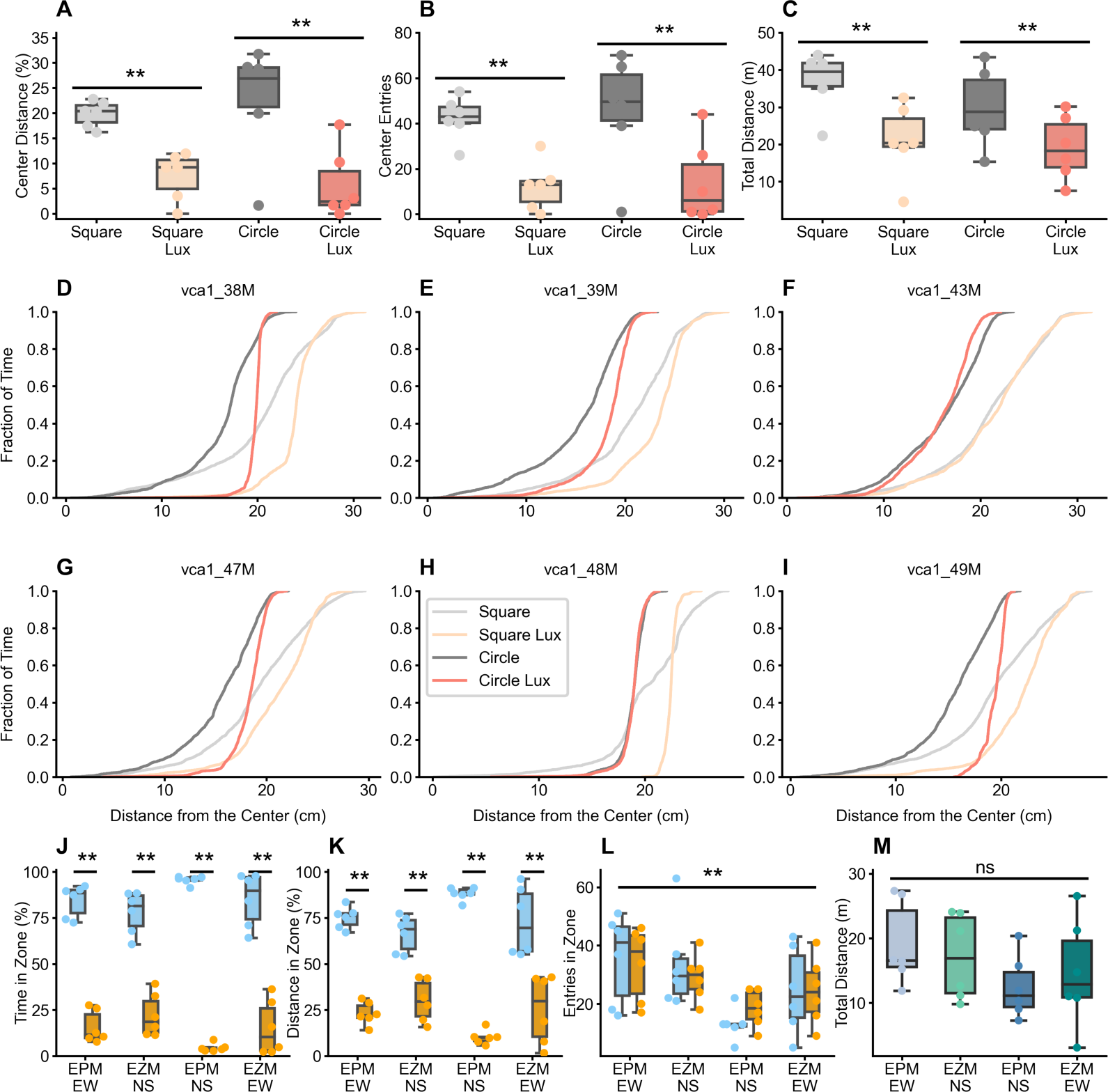
**(A)** The percent distance travelled in the center region of the open field arenas is significantly decreased during high lux sessions compared to low lux session (Two-way ANOVA significant main effect of lux, F_(1,20)_= 25.873, p < 0.01). **(B)** The number of entries in the center region of the open field arenas is significantly decreased during high lux sessions compared to low lux session (Two-way ANOVA significant main effect of lux, F_(1,20)_= 20.421, p < 0.01). **(C)** The total distance travelled in the open field arenas is significantly decreased during high lux sessions compared to low lux session (Two-way ANOVA significant main effect of lux, F_(1,20)_= 12.706, p < 0.01). **(D-I)** Cumulative distribution function (CDF) plots of distance from the center for individual mice. **(J)** The percent time in the open regions is significantly decreased compared to the closed regions in all sessions (Two-way ANOVA significant main effect of open/closed region, F_(1,40)_= 574.393, p < 0.01). **(K)** The percent distance travelled in the open regions is significantly decreased compared to the closed regions in all sessions (Two-way ANOVA significant main effect of open/closed region, F_(1,40)_= 271.906, p < 0.01). **(L)** The number of entries in the open and closed regions are significantly different across the elevated plus/zero maze sessions (Two-way ANOVA significant main effect of session, F_(3,40)_= 6.144, p < 0.01). **(M)** The total distance travelled is not significantly different across elevated plus/zero maze sessions (One-way ANOVA no significant effect of session, F_(1,20)_= 1.129, p = 0.36).

**Figure S2.**
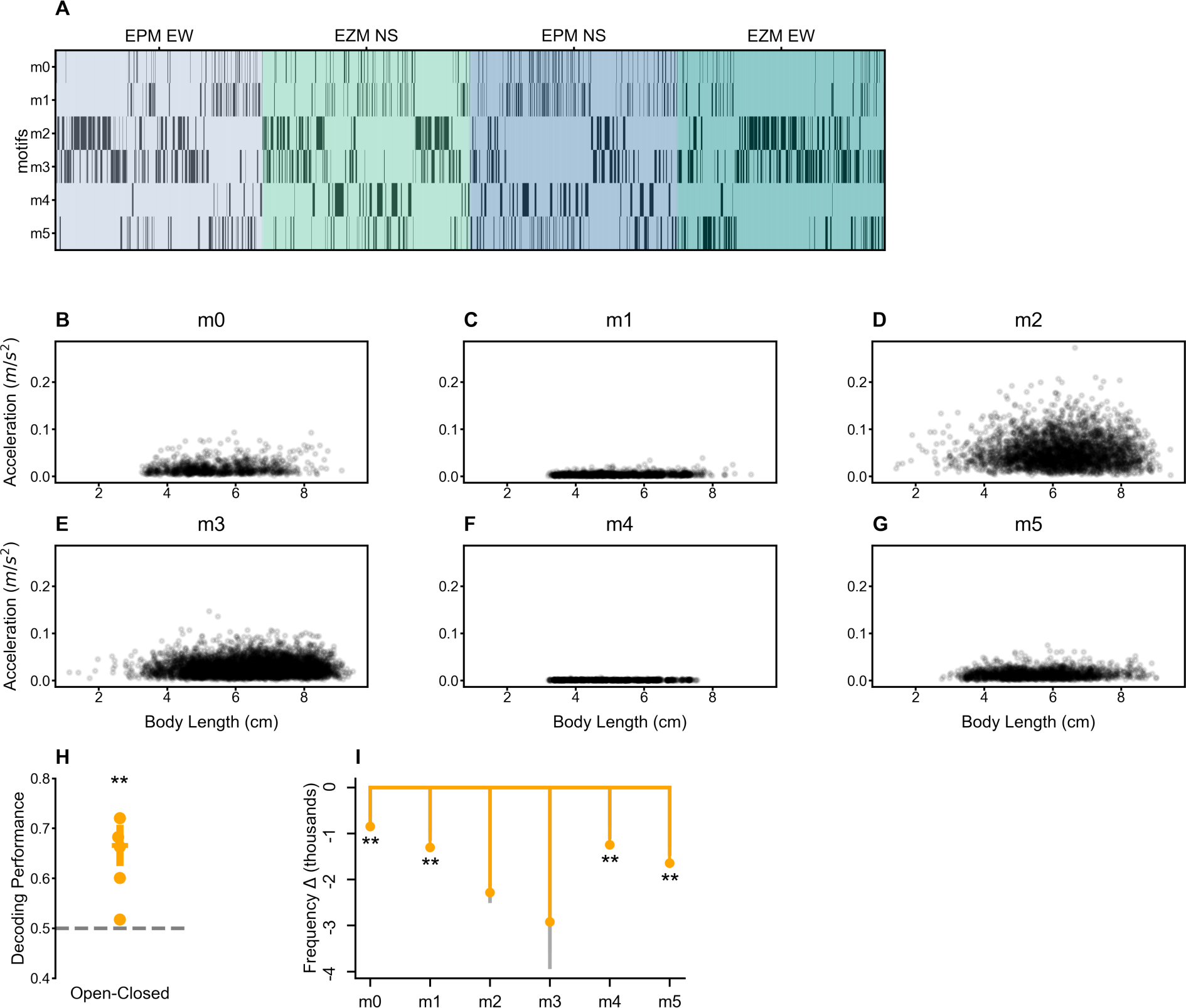
**(A)** An example motif raster plot showing the occurrence of each behavioral motif (m0 to m5) from a single mouse across the four elevated plus/zero maze sessions. **(B)** A scatterplot showing the relationship between body length and total head acceleration during motif m0. **(C)** A scatterplot showing the relationship between body length and total head acceleration during motif m1 for all mice. **(D)** A scatterplot showing the relationship between body length and total head acceleration during motif m2 for all mice. **(E)** A scatterplot showing the relationship between body length and total head acceleration during motif m3 for all mice. **(F)** A scatterplot showing the relationship between body length and total head acceleration during motif m4 for all mice. **(G)** A scatterplot showing the relationship between body length and total head acceleration during motif m5 for all mice. **(H)** A linear SVM trained on behavioral motifs can distinguish between open and closed regions (t-test, t_(5)_= 4.02, p < 0.05). **(I)** Motif frequencies of motifs m0, m1, m4, and m5 are significantly different between open and closed regions across all elevated plus/zero maze sessions. The observed frequency differences are represented by the colored bars and points. Grey bars represent the 95^th^ and 5^th^ percentile values of the null distribution. Asterisks represent the significance of motif frequency differences where * is p < 0.05 and ** is p < 0.01.

**Figure S3.**
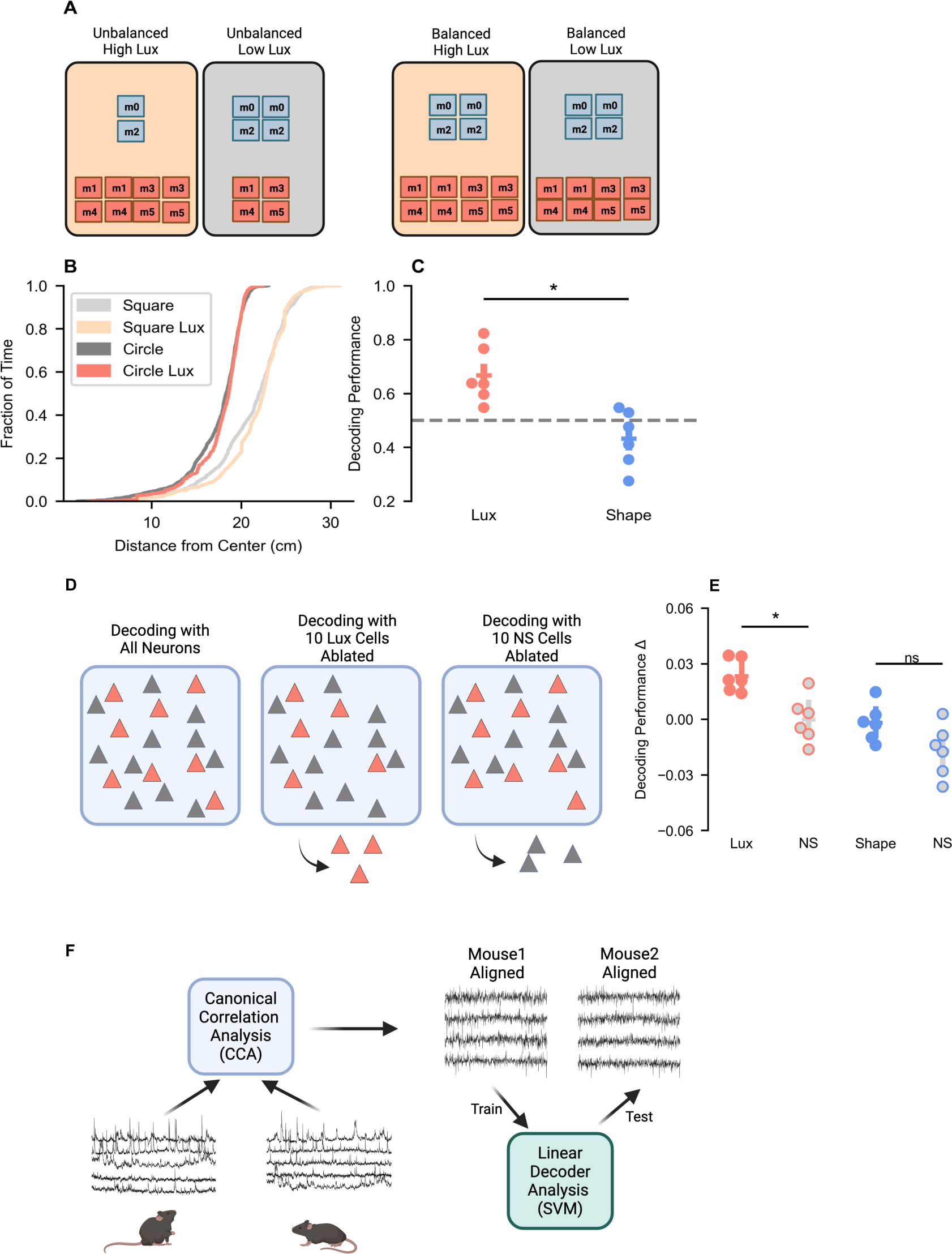
**(A)** Diagram depicting motif balancing procedure performed for decoding high vs. low lux and square vs. circle sessions from vCA1 neural data. **(B)** The distributions of distances from the center are similar in high and low lux conditions after the neural data has been manually balanced. **(C)** For manually balanced data, lux decoding performance is significantly above chance levels (paired t-test, t_(5)_= 4.76, p < 0.01); shape decoding performance is not significantly above chance levels (paired t-test, t_(5)_= −0.75, p = 0.49); lux decoding performance is significantly higher than shape decoding performance (paired t-test, t_(5)_= 3.22, p < 0.05). **(D)** Diagram depicting procedure performed for selective cell ablation decoding analysis of high vs. low lux and square vs. circle sessions. **(E)** The decoding performance drop is significantly greater when lux selective versus non-selective neurons are removed from the analysis (paired t-test, t_(5)_= 3.76, p < 0.05); the decoding performance drop is not significantly different when shape selective versus non-selective neurons are removed from the analysis (paired t-test, t_(5)_= 2.45, p = 0.06). **(F)** Diagram depicting procedure performed for aligning neural data from two different mice using canonical correlation analysis (CCA) and subsequent neural population decoding analysis.

**Figure S4.**
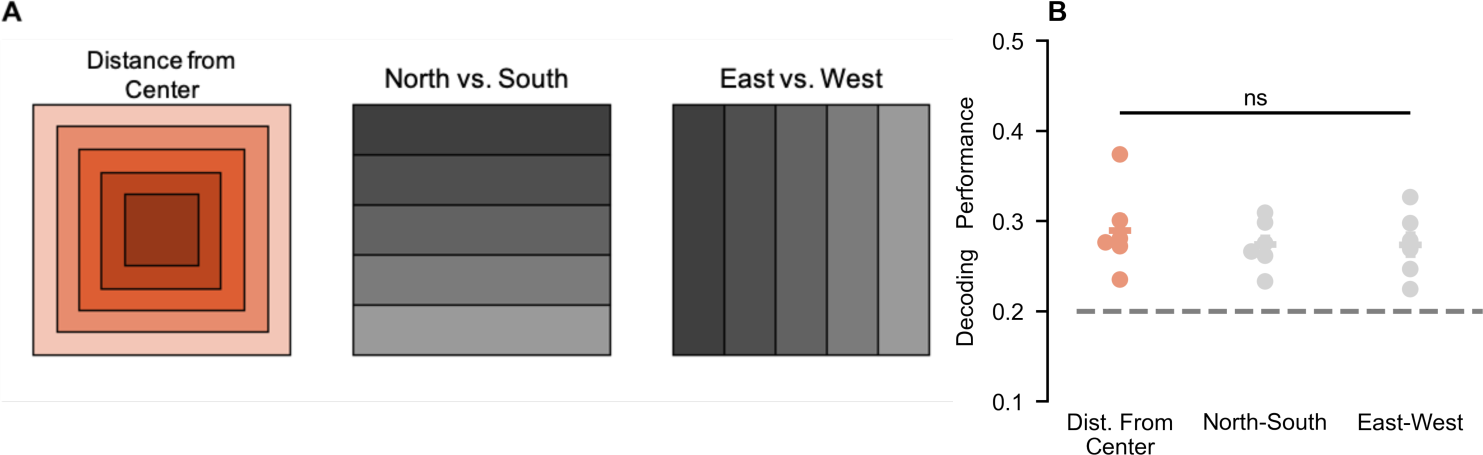
**(A)** Diagram depicting procedure performed for decoding distance from the center, north vs. south, and east vs. west. **(B)** Distance from the center, north-south, and east-west decoding performances are all significantly greater than chance (t-tests, t_(5)_= 4.74, p < 0.01, t_(5)_= 6.63, p < 0.01, t_(5)_= 4.99, p < 0.01), but they are not significantly different from each other (One-way ANOVA, F_(2,15)_= 0.36, p = 0.70).

**Figure S5.**
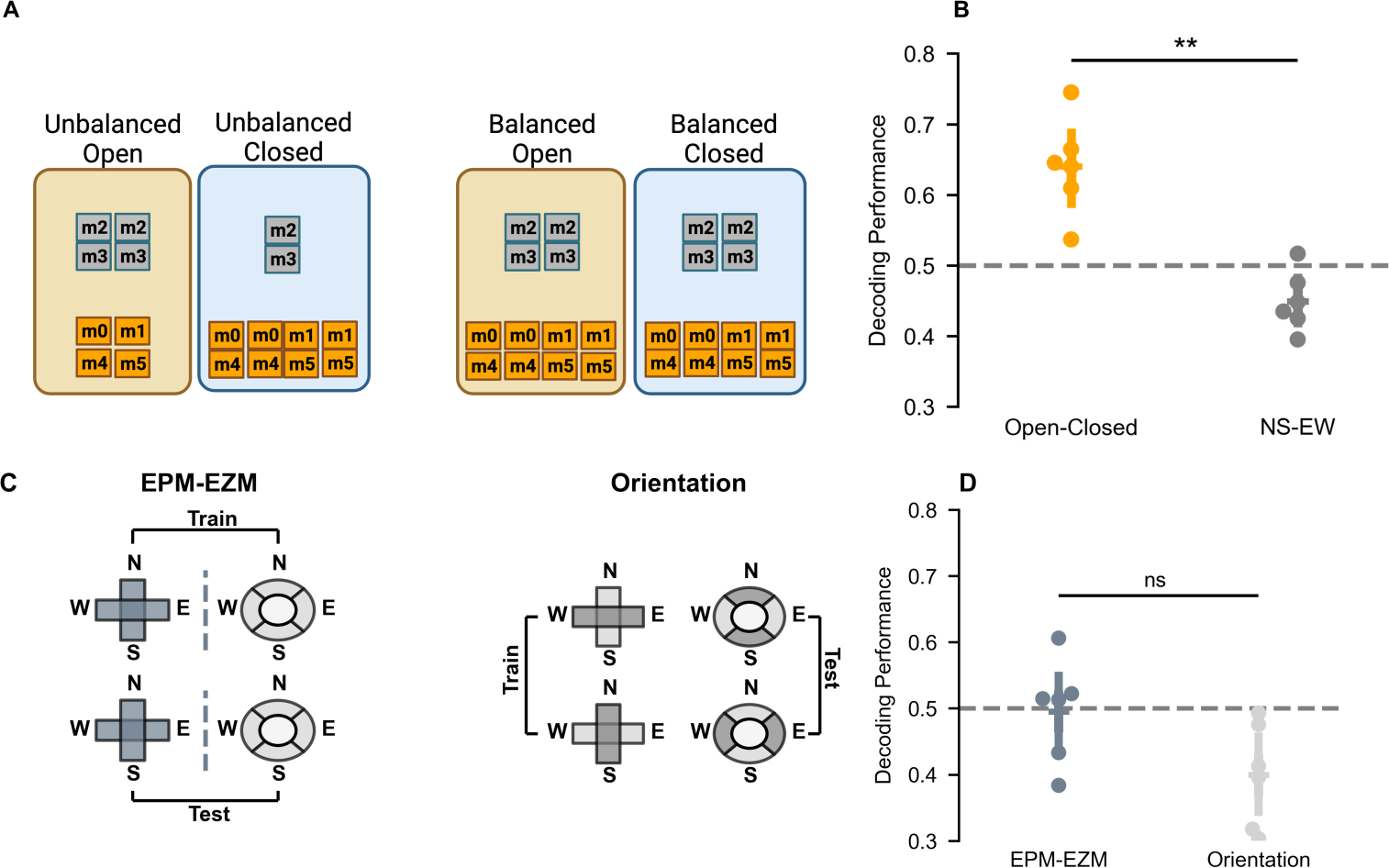
**(A)** Diagram depicting motif balancing procedure performed for decoding open vs. closed regions from vCA1 neural data. **(B)** For manually balanced data, open-closed decoding performance is significantly above chance levels (paired t-test, t_(5)_= 3.54, p < 0.05); NS-EW decoding performance is significantly below chance levels (paired t-test, t_(5)_= −2.06, p = 0.09); open-closed decoding performance is significantly higher than NS-EW decoding performance (paired t-test, t_(5)_= 4.26, p < 0.01). **(C)** Diagram depicting procedure performed for decoding elevated plus vs. elevated zero maze shapes and North-South vs. East-West orientations. **(D)** Shape decoding performance is not significantly different from chance levels (paired t-test, t_(5)_= 0.04, p = 0.97); orientation decoding performance is significantly lower than chance levels (paired t-test, t_(5)_= −3.23, p < 0.05); shape decoding performance is not significantly different than orientation decoding performance (paired t-test, t_(5)_= 1.86, p = 0.12).

**Figure S6.**
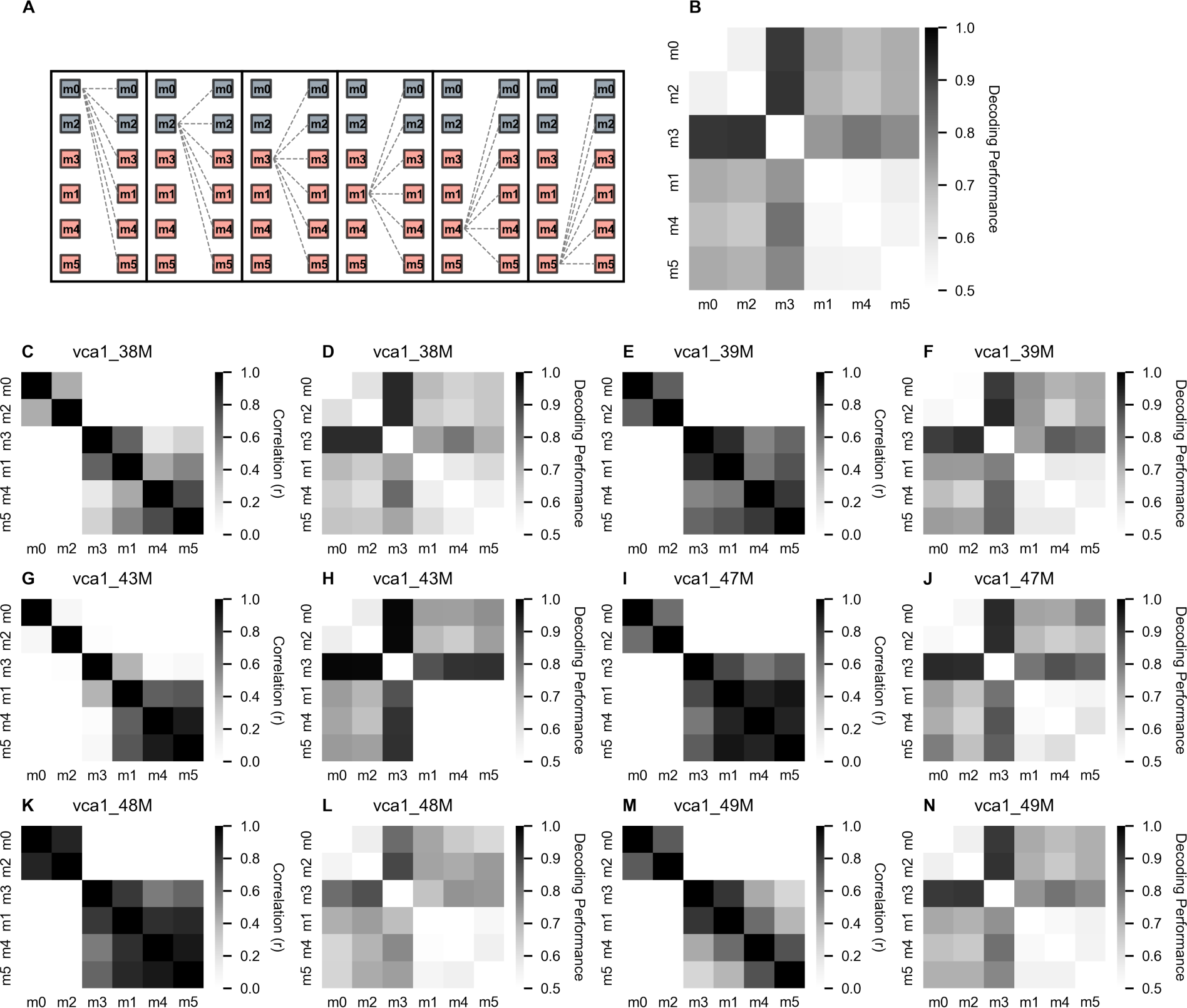
**(A)** Diagram depicting procedure performed for motif pair decoding analysis. **(B)** The average decoding performance for each pair of open field motifs across all mice. **(C, E, G, I, K, M)** Population vector correlation coefficient matrices for individual mice ordered to show the neural similarity between Cluster 1 and Cluster 2 motifs. **(D, F, H, J, L, N)** The average decoding performance for each pair of open field motifs for individual mice.

**Figure S7.**
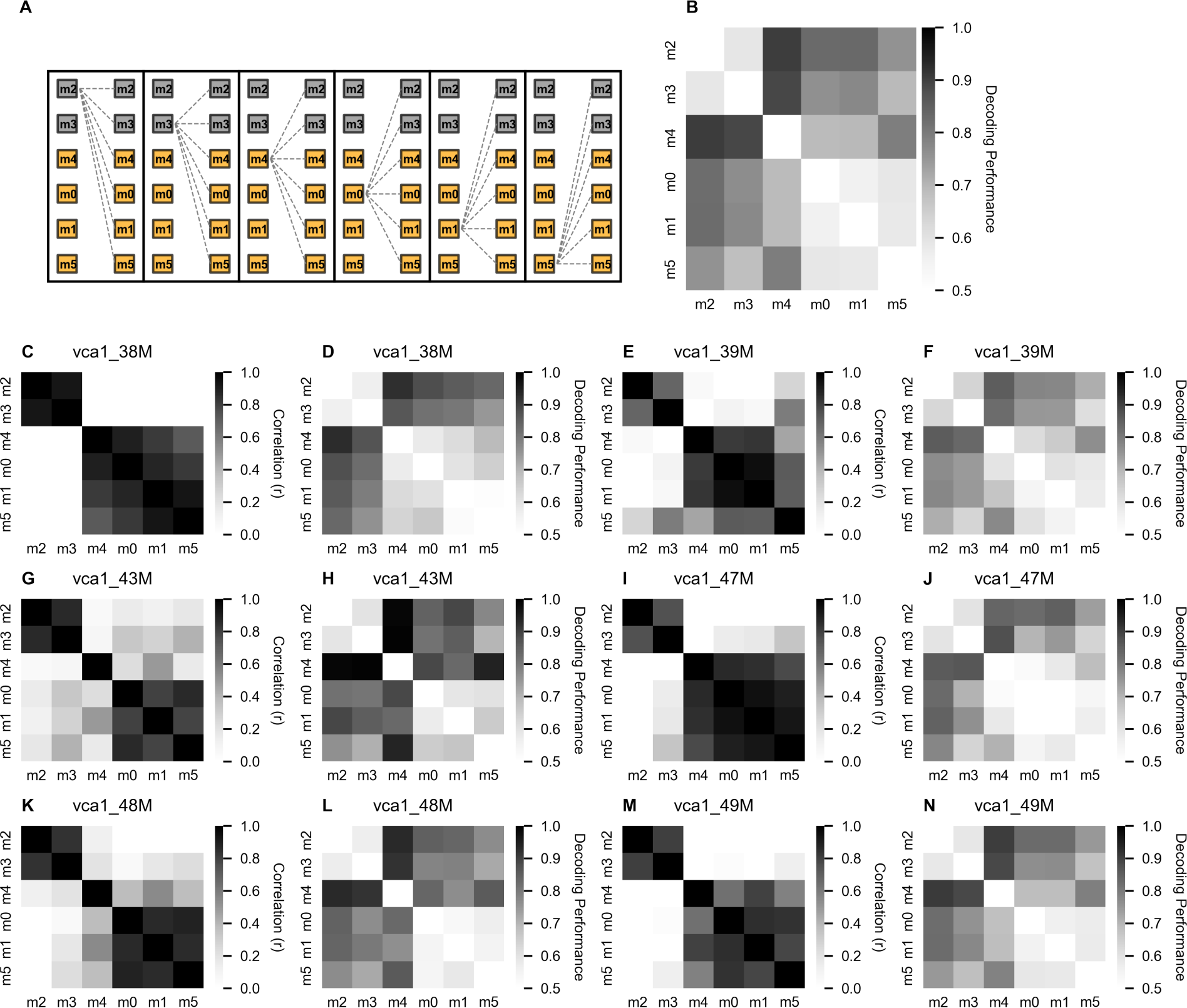
**(A)** Diagram depicting procedure performed for motif pair decoding analysis. **(B)** The average decoding performance for each pair of elevated plus/zero maze motifs across all mice. **(C, E, G, I, K, M)** Population vector correlation coefficient matrices for individual mice ordered to show the neural similarity between Cluster 1 and Cluster 2 motifs. **(D, F, H, J, L, N)** The average decoding performance for each pair of elevated plus/zero maze motifs for individual mice.

